# Paradox Found: Global Accounting of Lymphocyte Protein Synthesis

**DOI:** 10.1101/2022.10.31.514539

**Authors:** Mina O. Seedhom, Devin Dersh, Jaroslav Holly, Mariana Pavon-Eternod, Jiajie Wei, Matthew Angel, Lucas Shores, Alexandre David, Jefferson Santos, Heather D. Hickman, Jonathan W. Yewdell

## Abstract

Rapid lymphocyte cell division places enormous demands on the protein synthesis machinery. Flow cytometric measurement of puromycylated ribosome-associated nascent chains after treating cells or mice with translation initiation inhibitors reveals that ribosomes in resting lymphocytes *in vitro* and *in vivo* elongate at typical rates for mammalian cells. Intriguingly, elongation rates can be increased up to 30% by activation *in vivo* or fever temperature *in vitro*. Resting and activated lymphocytes possess abundant monosome populations, most of which actively translate *in vivo,* while *in vitro,* nearly all can be stalled prior to activation. Quantitating lymphocyte protein mass and ribosome count reveals a paradoxically high ratio of cellular protein to ribosomes insufficient to support their rapid *in vivo* division, suggesting that the activated lymphocyte proteome *in vivo* may be generated in an unusual manner. Our findings demonstrate the importance of a global understanding of protein synthesis in lymphocytes and other rapidly dividing immune cells.

## Introduction

Naïve lymphocytes are among the smallest nucleated cells in mammals – nearly devoid of cytoplasm, with few mitochondria – and have minimal metabolic activity, consistent with doubling times on the order of hundreds to thousands of days, respectively, for B cells ^1,2^ and T cells ^3^. Within a day of activation by cognate antigen, lymphocytes begin to divide rapidly, with reported doubling times as rapid as 6 hours ^4,5^. Such Jekyll and Hyde behavior requires massive induction of DNA and protein synthesis to support daughter cell production ^6^ as well as synthesizing large amounts of immune regulatory (e.g., cytokines) and effector molecules (*e.g.,* antibody and cytokines) ^7–9^.

Pioneering studies of protein synthesis regulation in lymphocytes utilizing radiolabeled amino acids on mitogen-activated human peripheral blood lymphocytes reported 7 to 20-fold increases in protein synthesis activity ^10,11^. While this is an impressive increase, it was assumed that sufficient protein was synthesized to enable the generation of daughter cells with the same protein content as their progenitor. Moreover, radiolabeling, like all methods, is imperfect, and its accuracy as a measure of protein synthesis rates depends on assumptions that are nearly impossible to definitively verify ^12^. Applying new methods to old problems is a tried-and-true method for generating new insights and discoveries.

Indeed, newer methods, including ribosome profiling ^13^, tRNA arrays ^14^, and tandem mass spectrometry ^15^, are revolutionizing the field of protein synthesis. This includes extending classical methods. Puromycin (PMY) is an aminonucleoside antibiotic that mimics tyrosine-tRNA, binding the ribosome A site and causing rapid chain termination by covalently attaching to the C-terminus of the nascent chain. PMY was first applied in classical protein synthesis studies ^16^ and remains a workhorse in understanding ribosomal catalysis of protein synthesis ^17–20^.

We developed the ribopuromycylation method (RPM) to better localize and quantify active protein synthesis. RPM uses a brief pulse of PMY to label elongating nascent chains frozen on ribosomes by treating cells with a translation elongation inhibitor. Ribosome-bound nascent chains are then detected using a PMY-specific monoclonal antibody in fixed and permeabilized cells via standard immunofluorescence ^21^ or flow cytometry ^22^.

Here, we use RPM, and the ribosome transit assay (RTA), an extension of RPM that measures elongation rates, in conjunction with classic techniques to quantify the number and protein synthesis activity of ribosomes in resting and activated human and mouse lymphocytes. Our findings reveal novel features of lymphocyte translation as well as a discrepancy in the protein synthesis capacity of T cells with respect to their rapid *in vivo* division rates, emphasizing the importance of quantitative accounting as a reality check for our limited understanding of fundamental aspects of cell biology and immunology.

## Results

### Characterizing protein synthesis in human lymphocytes *ex vivo* with flow RPM implicates widespread ribosome stalling in non-activated cells

We first used flow RPM to compare translation in non-activated *vs.* PMA/ionomycin/IL-2-activated human lymphocyte subsets after 2 and 5 days in culture (Figure 1A). We devolved the total flow RPM signals into T cell (CD4^+^, CD8^+^) and B cell (CD19^+^) subsets to follow distinct patterns of protein synthesis in each population (Supplemental Figure 1A). Comparing lymphocytes from 3 donors revealed considerable donor heterogeneity in RPM staining of day 2 activated cells and proliferation of lymphocyte subpopulations.

**Figure 1.**
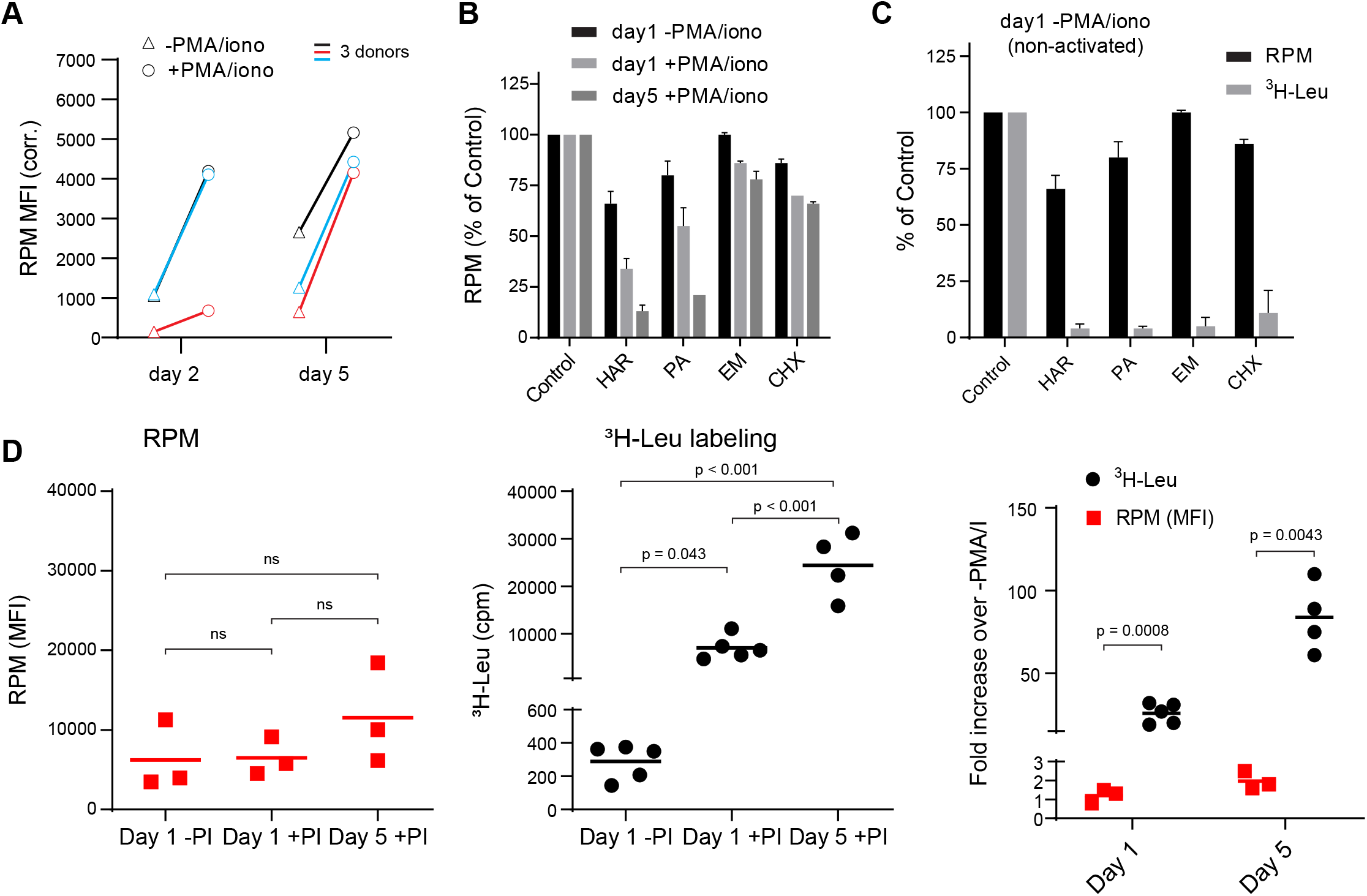
Stalled ribosomes in resting *ex vivo* human lymphocytes. (A) Primary human lymphocytes from three independent donors were cultured in PMA/ionomycin and IL-2 (+PMA/iono) or IL-2 only (-PMA/iono) for up to 5 days. CD45^+^ cells were processed for flow RPM. (B) Primary human lymphocytes were cultured *ex vivo* as indicated, followed by a 15 minute treatment with vehicle, harringtonine (HAR, 5μg/mL), pactamycin (PA, 10μM), emetine (EME, 25μg/mL), or cycloheximide (CHX, 200μg/ml), and all cultures were then treated with puromycin (PMY, 50µg/mL) for 5 minutes. Cells were harvested, and RPM staining was performed. Gated on CD45^+^ cells. Error bars represent standard deviation of two independent experiments. (C) Radioactive amino acid incorporation (0.2 mCi/mL [^3^H]-Leu for 5 min) or RPM (as in B) in day 1 non-activated human lymphocytes. Error bars represent standard deviation of two independent experiments. (D) Radioactive amino acid incorporation and RPM in rested and activated human lymphocytes. RPM MFI values (gated on CD45^+^ cells) on the left, [^3^H]-Leu incorporation (cpm) in the middle, and ratios of the activated to the resting cells on the right. Each point represents a single donor; bars indicate the mean from 3-5 independent donors. Left and middle panels: one-way ANOVA pairwise p-values; right panel: unpaired t-test p-values with Welch’s correction.

We performed RPM on peripheral blood mononuclear cells labeled with CFSE to track cell division by dye dilution (Supplemental Figure 1B). On day 2, activated CD8+ T cells demonstrated a wide range of RPM staining, with nearly all divided cells at day 5 CFSE^low^ and RPM^high^. Some divided cells exhibited near baseline RPM signals, however, consistent with their return to a resting state. Interestingly, although non-activated cells did not divide, ∼50% demonstrated increased RPM staining.

We noted that the RPM signal in PMA/ionomycin-activated CD8+ T cells was only 2-to 5-fold higher than in non-activated cells. This increase is modest compared to the ∼15-fold activation-induced increase in protein synthesis in original studies ^10,11^. To examine this discrepancy, we first incubated cells for 15 min with initiation inhibitors (harringtonine, HAR; pactamycin, PA) or elongation inhibitors (emetine, EME; cycloheximide, CHX), followed by RPM staining. Elongation inhibitors had minor effects on RPM of activated or resting cells (Figure 1B), as expected due to ribosome retention of nascent chains ^21^. Initiation inhibitors, however, clearly discriminated between resting and activated cells. RPM signal was diminished by up to 80-90% on day 5 post-activation. Note that at the standard translation rate of 6 amino acids/sec, 15 min is sufficient time to complete translation of all but the very longest transcripts.

We repeated this experiment using day 1 resting lymphocytes to directly compare flow RPM with classical metabolic radiolabeling with [^3^H]-Leucine ([^3^H]-Leu) (Figure 1C). All inhibitors nearly completely blocked incorporation of [^3^H]-Leu into proteins, suggesting that there were actively translating ribosomes in resting cells and that the inhibitors were active, even though RPM labeling was only weakly impacted. We also performed a time course examining [^3^H]-Leu incorporation compared to flow RPM signal in day 1 resting, day 1 activated, and day 5 activated human lymphocytes. Plotting the ratios between activated and non-activated cells from RPM flow vs. [^3^H]-Leu incorporation revealed a substantial difference between the two methods (Figure 1D).

Thus, we cannot attribute the persistence of flow RPM staining in translation initiation inhibitor-treated resting lymphocytes to incomplete inhibition of protein synthesis. Instead, these data are consistent with a significant fraction of “stalled” ribosomes in cultured resting cells, *i.e.,* ribosomes with nascent chains that are not actively translating. Stalled ribosomes would be labeled with PMY, as originally described in neurons ^23^, but would not incorporate [^3^H]-Leu, just as we observe with resting lymphocytes.

### Flow RPM measures ribosome elongation rates in live cells

To extend these findings, we developed a variation of approaches that use initiation inhibitors to measure ribosome transit times, for example by conversion of polysomes to monosomes ^24^ or ribosome profiling ^13^. To derive a relative ribosome transit rate, we incubate cells with the initiation inhibitor HAR for increasing times before shifting cells to 4°C to halt ribosome elongation and process for RPM staining (Figure 2A).

**Figure 2.**
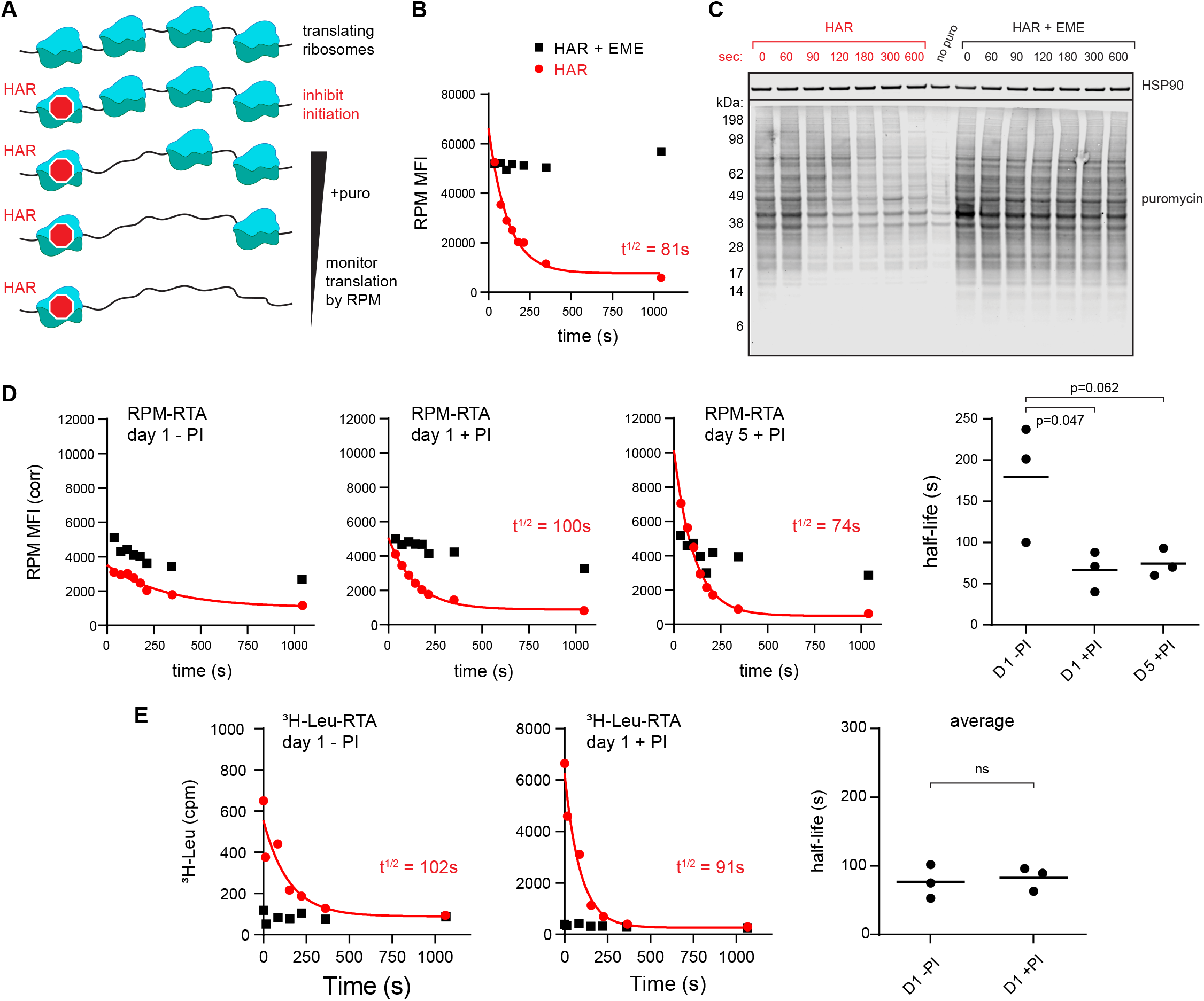
RPM measures ribosome transit times in HeLa and human lymphocytes. (A) Schematic representation of the ribopuromycylation (RPM) Ribosome Transit Analysis (RTA) method. Translation initiation is blocked and the decrease in RPM is monitored as the elongating ribosomes run off mRNA. (B) RPM-RTA in HeLa cells. Harringtonine (HAR, 5μg/mL) is used to inhibit new ribosome initiation; emetine (EME, 25μg/mL) is used to freeze ribosomes on mRNA; puromycin (PMY, 50μg/mL) generates RPM signal. Curve is fitted using one phase exponential decay, and ribosome transit times are expressed as RPM half-time to decay. (C) Same as B, but cells are instead lysed in the presence of MG-132 and subjected to anti-puromycin western blot analysis. (D) Representative plots of the RPM-RTA signal in resting and activated human lymphocytes (left three panels). Gated on CD45+ cells. Far right, ribosome transit times determined from 3 independent donors. Each dot represents data from one individual donor; the horizontal bars indicate the mean. P-values indicate one-way ANOVA pairwise comparisons. (E) Ribosome transit times as in A but determined by [^3^H]-Leu incorporation instead of RPM. After treatment with HAR or HAR plus EME, cells were labeled for 5 minutes in 0.25mCi/mL [^3^H]-Leu. Right panel, ribosome transit times determined by [^3^H]-Leu incorporation from three independent donors. Each dot represents data from one individual donor; the horizontal bars indicate the mean. Unpaired t-test.

We validated this approach in HeLa cells whose ribosome transit times are well characterized ^25^. This revealed a curve that follows one phase exponential decay (Figure 2B; gating strategy in Supplemental Figure 1C), with a calculated half-life to decay of 70-150 seconds. Including EME with HAR prevented decay of the RPM signal, as predicted, since EME blocks elongation while enabling (even enhancing) puromycylation ^21,26^.

Immunoblotting of puromycylated nascent chains validated the approach by showing a time-dependent decrease in puromycin signal and increased Mr of nascent chains after blocking initiation (Figure 2C). This is expected since nascent chains present at later time points after blocking initiation will be longer. Incubation of EME with HAR greatly retarded the loss of signal and the shift to longer puromycylated nascent chains.

We applied this RPM-based “ribosome transit assay” (RTA) to investigate translational control in human lymphocytes. Day 5 activated lymphocytes behaved similarly to HeLa cells in their RTA half-life and EME sensitivity (Figure 2D). By contrast, in day 1 resting lymphocytes, there was a limited decay in the signal. Further, the decay was similar in EME-treated cells, consistent with the idea that the flow RPM signal in day 1 resting lymphocytes predominantly represents stalled ribosomes with bound nascent chains.

To independently measure ribosome transit times in day 1 resting *vs.* activated lymphocytes, we treated cells for increasing times with HAR and then pulse-labeled with [^3^H]-Leu (Figure 2E). This showed that both resting and activated cells demonstrated a decay half-life of ∼90-100 seconds, similar to the RTA values for activated lymphocytes and HeLa cells.

Based on these findings, we conclude that:

1. A large fraction of ribosomes stalled in resting cultured lymphocytes.
2. Elongation occurs at similar rates for HeLa cells and lymphocytes, with the active ribosomes in resting lymphocytes translating at a similar rate as fully activated lymphocytes.
3. RTA provides a simple flow cytometric measure of ribosome transit rates, confirming and extending the findings of Arguello *et al.* ^27^ who reported a highly similar method.

### Resting human lymphocytes have a dominant monosome population

Protein synthesis is generally believed to occur predominantly in polysome structures, consisting of multiple ribosomes transiting a single mRNA ^28^. Classic ^29,30^ and more recent studies ^31^ have established, however, that resting lymphocytes have few polysomes and provided evidence for active monosome translation by their stability in high salt, which dissociates non-translating ribosomes ^32^.

Confirming these reports, we found that a large fraction of assembled ribosomes in resting human lymphocytes fractionate as monosomes in sucrose gradients (Supplemental Figure 2A). Polysome abundance increases over two days post-activation. Treating freshly isolated human lymphocytes with CHX to freeze ribosomes ^33,34^ did not increase polysome recovery (Supplemental Figure 2B). These findings, coupled with our RPM/RTA measurements, indicate that stalled ribosomes are likely monosomes.

### Protein synthesis in mouse lymphocytes *ex vivo*

Working with human lymphocytes is problematic – preparations between individuals vary considerably, and the manipulations required to isolate lymphocytes from donor blood, such as elutriation and Percoll gradient purification, increase the time cells spend outside their physiological environment.

Seeking a more reproducible system without the impact of potential artefactual stalling of the translation machinery, we turned to OT-I TCR transgenic mice ^35^. OT-I cells are CD8^+^ T cells specific for a defined cognate ligand (mouse K^b^ MHC class I molecules bound to the ovalbumin-derived SIINFEKL peptide) that can be activated *in vitro* or *in vivo*. OT-I T cells can be obtained from spleen or lymph nodes (LNs) in reasonable numbers at ∼80% purity without further manipulation.

RTA analysis revealed that there was no decay in RPM signal for *ex vivo* day 1 resting OT-I T cells, consistent with near total stalling of once-translating ribosomes, as we identified in human lymphocytes (Figure 3A, middle panel). By contrast, in freshly isolated OT-I cells, the RPM signal decays by 50%, consistent with active translation by 50% of the ribosomes with the rest of the ribosomes likely stalled or poised. The signal decay t½ of 40 seconds is consistent with translation of shorter than average mRNAs or stalling on partially translated mRNAs. Polysomes were a minor fraction in freshly isolated mouse lymphocytes, even when mice were pre-treated with CHX (Supplemental Figure 2C, further addressed in the next section).

**Figure 3.**
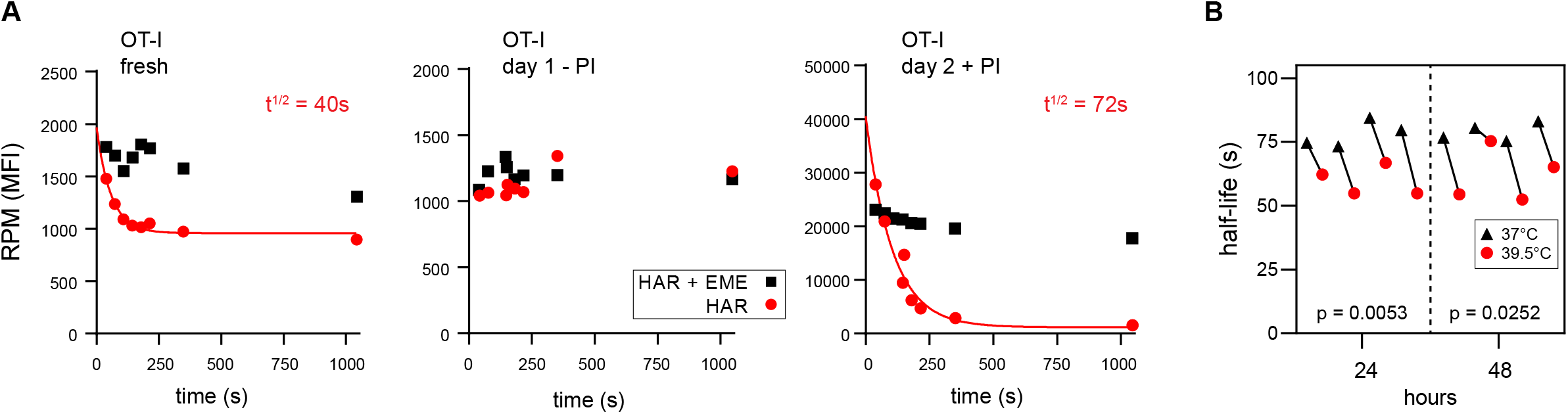
RPM ribosome transit analysis of OT-I T cells *in vitro*. (A) Lymphocytes from spleens and lymph nodes from transgenic OT-I mice were isolated, and either used immediately, cultured for one day in the absence of PMA/ionomycin, or cultured for 2 days in the presence of PMA/ionomycin and IL-2. RPM-RTA analysis was conducted to determine ribosome transit half-lives, both with and without EME. (B) Lymphocytes from spleens and lymph nodes from transgenic OT-I mice were isolated, labeled with CFSE, and cultured under activating conditions for either 24 or 48 hours. Cells were harvested, and RPM-RTA was performed at both 37°C and 39.5 °C. Half-life of RPM signal by RTA is plotted; p-values determined by paired t-test analysis.

By contrast, day 2 activated *ex vivo* OT-I T cells demonstrated a 20-fold increased RPM signal relative to resting cells, a near total signal decay with a t½ of ∼70 seconds (Figure 3A, right panel), and a preponderance of polysomes (Supplemental Figure 2D). This is consistent with the large fractional engagement of ribosomes upon activation. Notably, the decay rate is faster than observed in previous conditions and intriguingly, the rate increases by ∼20% at a “fever” temperature of 39.5 C° (Figure 3B). This suggests that lymphocytes may be able to exceed the standard mammalian cell elongation rate of ∼6 residues/second ^36^, particularly under fever conditions, when maximizing T cell protein synthesis is likely at a premium to support their anti-viral activity by rapid division and production of effector molecules.

### Protein synthesis in mouse lymphocytes and innate immune cells *in vivo*

Mammalian cells evolved, of course, in mammals, not in plastic flasks nurtured by synthetic media in a 20% oxygen atmosphere. We therefore adapted the RTA assay to mice. To simultaneously measure resting and activated T cells, we adoptively transferred CFSE-labeled OT-I T cells into congenic B6 mice, which we infected with SIINFEKL-expressing vaccinia virus (VACV) to activate OT-I cells. We then injected mice with HAR for 0-10 min, followed by PMY injection and flow RPM processing of harvested splenocytes (Figure 4A). With each mouse providing a single data point, we could generate RTA curves for non-activated host CD4 and CD8 cells as well as transferred OT-I cells activated by VACV infection (Figure 4B). These curves show that nearly all ribosomes with nascent chains in both resting and activated lymphocytes are actively elongating proteins *in vivo*.

**Figure 4.**
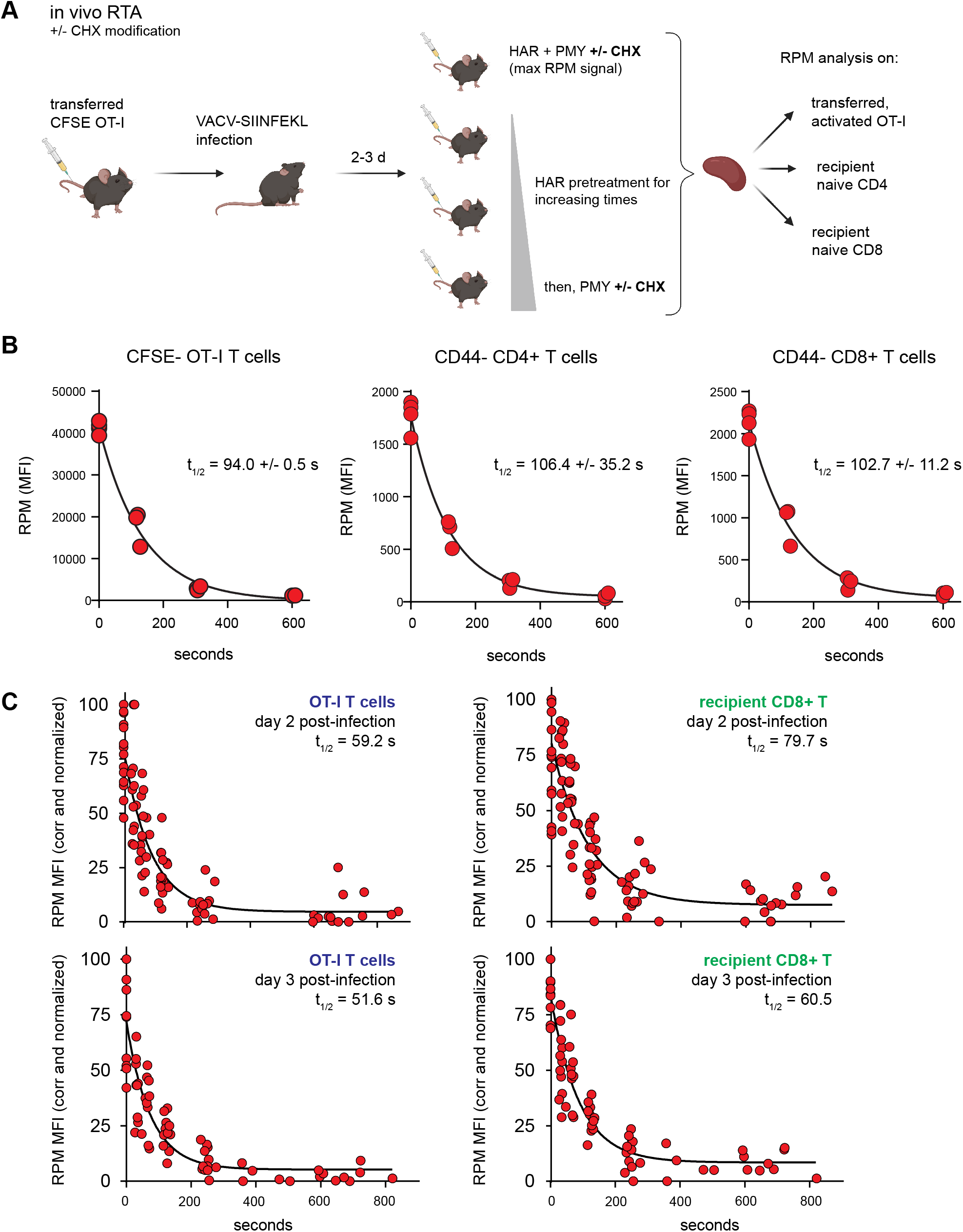
Translation rates of resting and activated T cells *in vivo*. (A) Depiction of the *in vivo* RPM-RTA method. Labeled OT-I T cells are first adoptively transferred, followed by VACV-SIINFEKL infection of mice. RTA analysis is performed by intravenous injection of HAR followed by PMY (+/- CHX to prevent leakiness from HAR inhibition alone). Spleens are harvested for RPM analysis on both endogenous and transferred T cells. Schematic designed with Biorender. (B) CFSE-labeled Ly5.2^+^ (CD45.2^+^CD45.1^-^) OT-I T cells were adoptively transferred into Ly5.1 (CD45.1^+^CD45.2^-^) mice, which were then infected with VACV-SIINFEKL to activate the OT-I cells. Three days after infection, mice were intravenously injected with HAR simultaneously with PMY for 5 minutes (maximum signal), or first injected with HAR for ∼110, ∼275, or ∼575 seconds before being injected with PMY for 5 minutes. Splenocytes from mice were harvested, surface stained for gating and activation markers as indicated, fixed and permeabilized, and stained for RPM. Gates were CFSE^low^ OT-I CD8^+^ T cells to measure decay in activated cells, and CD44^-^CD8^+^ or CD44^-^CD4^+^ T cells to measure decay in resting T cells. The curve was generated by fitting to a one phase exponential decay. Representative of two independent experiments, 2-4 mice per group, with the mean and standard deviation of the calculated half-life decays as indicated. (C) RTA, with the CHX modification, of adoptively transferred OT-I T cells or un-activated host CD8^+^ T cells in mice infected for 2 or 3 days with VACV-SIINFEKL. 3-4 independent experiments combined, normalized by setting maximum background-subtracted signal to 100.

The elongation rate *in vivo* is surprisingly slower than the *in vitro* rate. Notably, this experiment used our original protocol of PMY treatment alone ^22^ since EME, the inhibitor used to stabilize puromycylated polypeptides on ribosomes *in vitro* ^21^ was ineffective *in vivo*. We found, however, that CHX is active *in vivo*, arresting the accumulation of puromycylated polypeptides for at least 60 min after injecting PMY (Supplemental Figure 3A). We therefore modified the RTA by simultaneously treating animals with CHX with PMY to determine the relative amount of ribosome-associated nascent chains *in vivo*. This enabled comparison of translation activity in various immune cell types using 15 min HAR pretreatment values to subtract the signal from stalled ribosomes. The number of translating ribosomes varies over a narrow range among resting splenic lymphocytes, NK cells, macrophages, and neutrophils (Supplemental Figure 3B).

Using this improved RPM protocol, one day after infecting mice with VACV we now measured a ∼15-fold increase in translating ribosomes in activated OT-I T cells *in vivo* (Supplemental Figure 3C; gating strategy in Supplemental Figure 3D) as compared to the 10-fold increase we previously reported ^22^. As cell division progressed over the next two days, the signal from translating ribosomes decreased (Supplemental Figure 3E-F). Comparing the results from Supplemental Figure 3C (OT-I cells) with Supplemental Figure 1B (polyclonal human CD8+ T cells) reveals what we had described previously ^22^, that a transgenic T cell population has much less spread of RPM staining when compared to activated polyclonal T cells in C57/BL6J mice after VACV infection, or here when comparing to all human CD8+ T cells.

We next performed the modified RTA to measure translation rates in OT-I cells *in vivo* on day 2 and 3 post-infection with VACV-SIINFEKL (Figure 4C). Addition of CHX to the *in vivo* RTA is important because of the well-characterized “leakiness” of harringtonine ^37^; indeed, ribosome transit times in activated OT-I cells were now in line with the *in vitro* rates, and were ∼20% faster than transit times in recipient (non-activated) T cells.

These results indicate that:

1. Monitoring accurate translation and rates *in vivo* is possible and avoids artifacts associated with *ex vivo* lymphocyte cultures.
2. Activation increases elongation rates in lymphocytes by ∼20%.

### Contribution of monosomes vs. polysomes to T cell translation

We next biochemically characterized translation in resting OT-I cells *in vivo* or OT-I cells activated *in vitro* by PMA/ionomycin/IL-2. We treated animals/cells with CHX/PMY, isolated ribosomes from cell lysates on sucrose gradients in monosome and polysome fractions, blotted fractions onto nitrocellulose and stained with antibodies against RPL7 or PMY. The robust PMY signal shows that, contrary to recent claims ^38,39^, PMY does not completely release nascent chains when ribosomes are previously exposed to CHX (Figure 5A).

**Figure 5.**
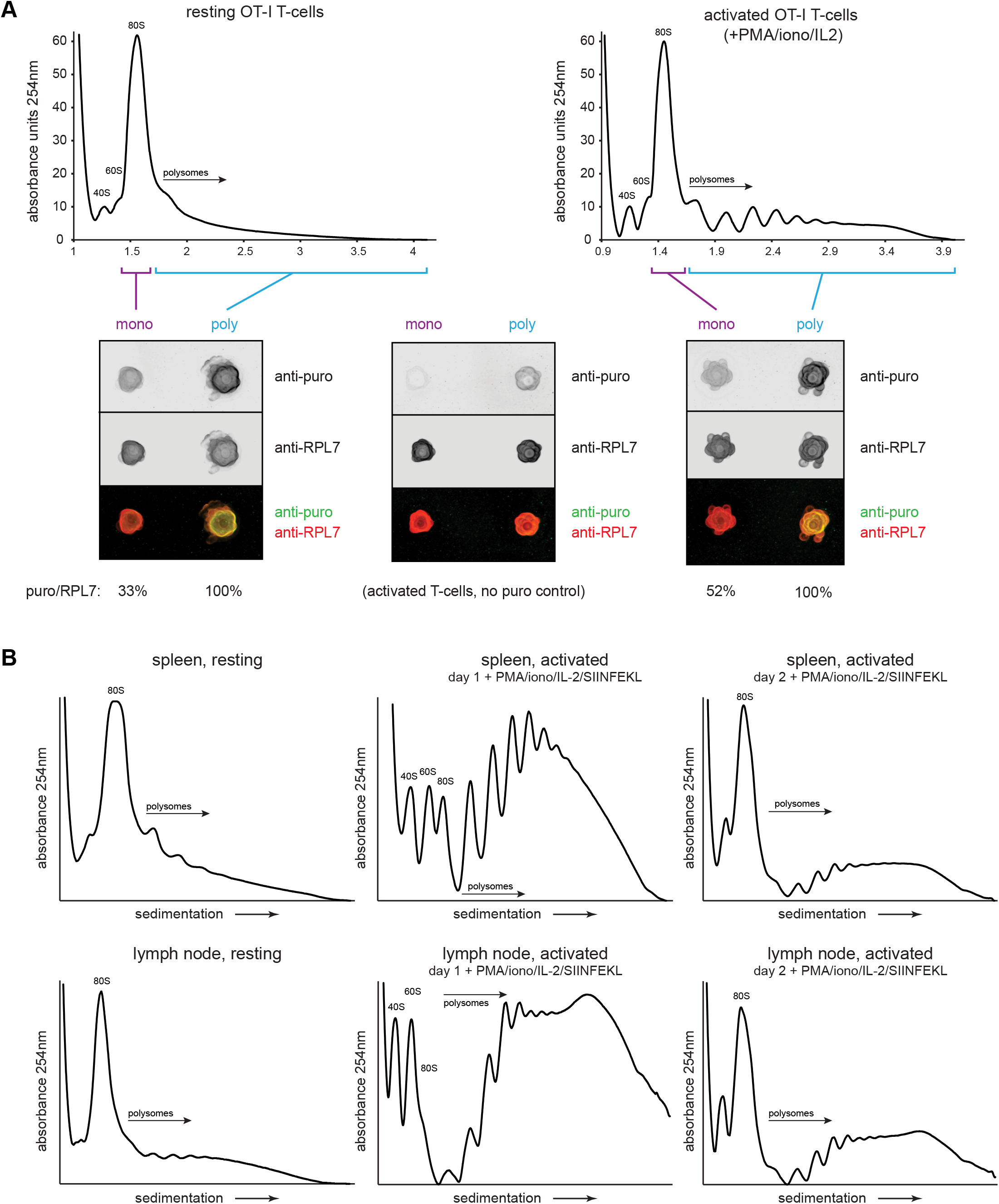
Puromycylation reveals percentage of actively translating monosomes in resting and activated T cells. (A) OT-I mice were treated intravenously with CHX and PMY, and lymphocytes from the spleens and lymph nodes were isolated and subjected to polysome profiling by ultracentrifugation through 15-45% sucrose gradients (resting OT-I T cells). OT-I T cells activated *in vitro* for 2 days with PMA/ionomycin and IL-2 (without cognate SIINFEKL peptide) were treated either with CHX alone (no PMY control) or CHX with PMY and subjected to polysome profiling. The indicated fractions were collected, pooled, and their ribosomes were re-isolated and dotted onto a nitrocellulose membrane for blotting with antibodies against PMY and RPL7. After subtraction of background signal from the anti-puro antibody (middle panel), the PMY/RPL7 ratio of monosomes was expressed relative to that of polysomes, which was defined as 100% translating. Representative of two independent experiments. (B) For resting T cells, OT-I mice were treated intravenously with CHX, and lymphocytes from the spleens or lymph nodes were isolated and lysed. For activated T cells, lymph node or splenic OT-I T cells were stimulated *in vitro* for 2 days with PMA/ionomycin, IL-2, and exogenous SIINFEKL, followed by treatment with CHX for 5 minutes. For both resting and activated cells, ribosome-containing lysates were fractionated via ultracentrifugation on 15-45% sucrose gradients.

After setting the puromycylation:RPL7 ratio in polysomes to 100% (assuming that all ribosomes in the polysome fraction are actively translating), we found that 33% of monosomes in resting *in vivo* OT-I T cells and 52% of monosomes in day 2 activated OT-I T cells were puromycylated. Since *in vivo* RTA indicates that there is essentially no stalling of puromycylated ribosomes (Figure 4C), these data demonstrate robust translation in T cell monosomes. Assuming equal elongation rates, ∼38% and ∼32% of overall translation would occur in monosomes of resting *in vivo* and activated *in vitro* cells, respectively. We note, however, that since PMY reduces the number of polysomes recovered from CHX-treated cells by 5-10%, a small fraction of translating monosomes probably derive from the polysome population.

The high fraction of monosome-based translation is surprising in activated cells. We noted that the activation protocol for OT-I T cells we used is far less effective than that published by Tan et al. ^31^ which includes SIINFEKL antigenic stimulation along with PHA and ionomycin. Bulk peptide-antigen stimulation directly *ex vivo* is not possible with human cells, but it is with transgenic murine T cells, and the methodological adaptation enhances activation. Direct comparison of the protocols confirmed the superiority of Tan *et al.*, as shown by activation markers (CD69, CD25, CD44), cell size (measured by side scatter, which correlates well with automated diameter measurements), and cell division (Supplemental Figure 4A-B). OT-I cells activated by this protocol yielded large increases in the observed polysome fractions of activated splenocytes, or lymph-node-derived lymphocytes (Figure 5B). This was also evident during high salt fractionation conditions, where we found that 500mM was necessary to fully dissociate non-translating ribosomes compared to the often used 300mM (Supplemental Figure 5A-B). T cell ribosomes had quantifiable but low levels of monosomes under these high salt conditions (Supplemental Figure 5C-D; note that we could not obtain enough activated OT-I cells *in vivo* for these experiments).

These findings indicate that monosomes make a major contribution to translation in resting T cells but are likely to make a minor contribution in fully activated cells. These results might also complicate the conclusion reached by Geraschchenko *et al.* ^40^ that harringtonine may only be useful until 45 seconds after the start of treatment, as the assumption was made that polysomes were the only ribosome subset actively translating mRNA.

### Accounting for translation in lymphocytes: measuring the protein-to-ribosome ratio

Cells need to synthesize sufficient proteins to regenerate a complete proteome each division cycle. This number will depend on the division rate, cell size, protein concentration, and protein loss due to degradation and export (secretion, release of exosomes, loss of other cellular material). To understand how the protein synthesis apparatus enables such rapid T cell division times, we quantitated a number of critical protein synthesis parameters in resting and activated OT-I T cells (Figure 6A). For these experiments, we used the optimized protocol for *in vitro* OT-I T cell activation ^31^ that greatly increased the fraction of ribosomes in polysomes on day 1 post activation.

**Figure 6.**
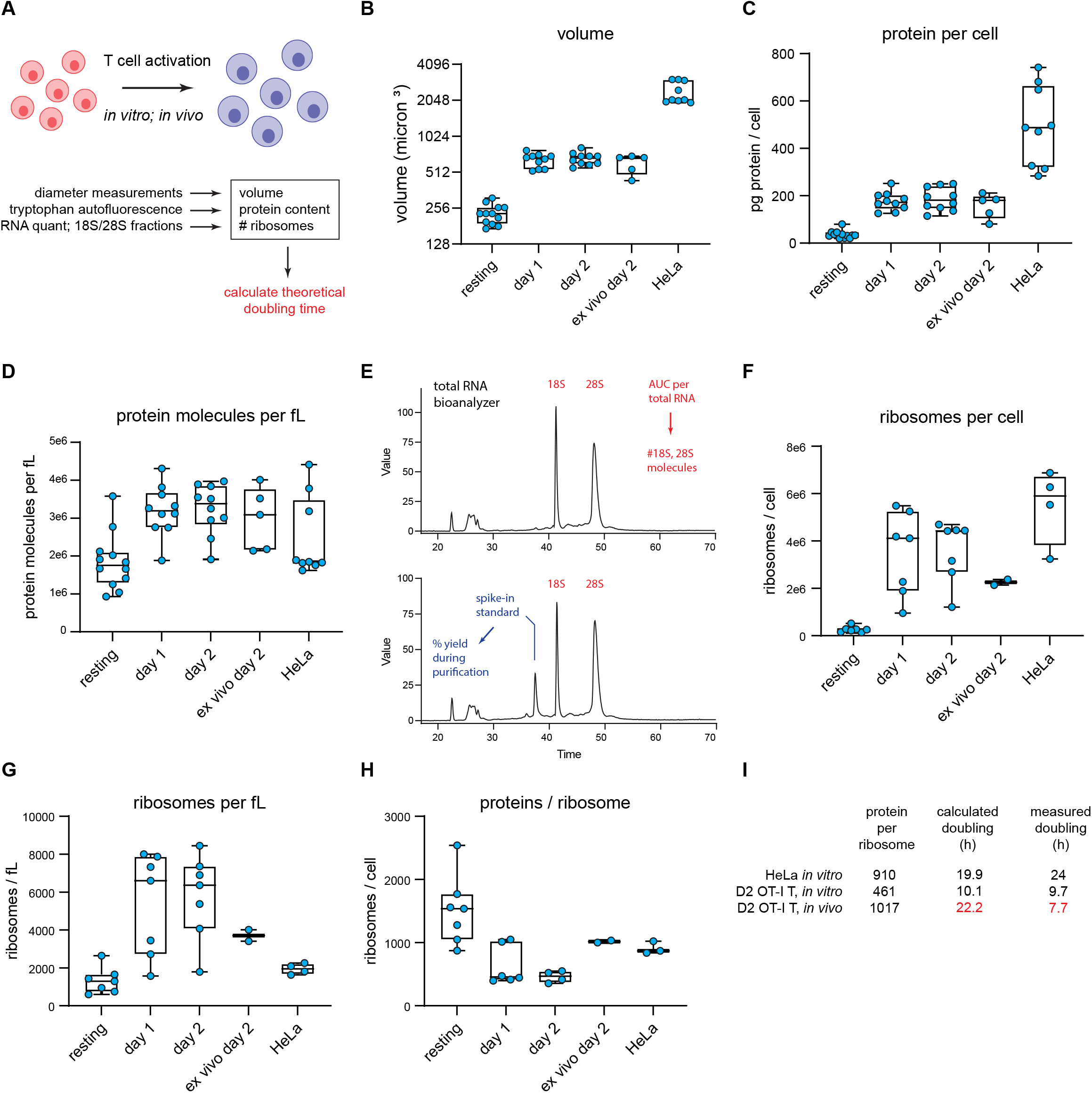
T cell accounting reveals discrepancy in proteome duplication rate for activated T cells *in vivo*. (A) Measurements made to calculate *in vitro* and *in vivo* rates of T cell division. (B) Volume calculations based on diameter measurements made by automated cell counter for the indicated cell types. Day 1 and day 2 represent *in vitro* activated OT-I T cells. *Ex vivo* day 2 represent cells activated *in vivo* for 2 days, followed by isolation and processing. (C) Protein content per cell as measured by tryptophan fluorescence of denatured lysates. (D) Protein molecules per fL, assuming an average protein length of 472 aa and average amino acid mass of 110 Da. (E) Example output from custom bioanalyzer method to determine number or ribosomes per cell. Total RNA is quantified and the bioanalyzer is used to determine area under the curve for 18S and 28S percentage of total RNA. Additionally, an exogenous mRNA standard is spiked into the sample prior to RNA isolation to determine the percent loss in yield during the purification procedure. Combined, this method allows for the accurate determination of total number of 18S and 28S molecules per cell. (F) Number of ribosomes per cell for the indicated cells. (G) Ribosome per fL for the indicated cells. (H) The protein/ribosome ratio, a representation of how many proteins a single ribosome would need to create to duplicate the proteome. (I) Discrepancy between measured and calculated rates of division for OT-I T cells activated and dividing *in vivo*.

Automated microscope measurements revealed that OT-I T cells increase in diameter from the resting state to the day 1 and day 2 activated states, with a corresponding calculated volume increase (based on spherical geometry) of ∼2.9 fold (Figure 6B). To quantify protein content, we determined total tryptophan (Trp) autofluorescence of fully denatured proteins in a total cell lysate ^41^. Protein content per T cell increases ∼5-fold following activation (Figure 6C), from 421 million proteins per cell (assuming an average length of 472 aa and a proteome Trp content of 0.69% ^41^) to 2.15 billion proteins per cell in day 2 activated cells, resulting in a net 1.7-fold increase in protein concentration (Figure 6D).

We determined the number of ribosomes per cell using a Bioanalyzer electrophoresis device to measure the amount of 18S and 28S rRNA in purified total RNA based on staining with a RNA-binding dye and utilizing a spike-in standard mRNA to control for yield loss during RNA purification (Figure 6E). The maximal number of translating ribosomes is limited by the less abundant subunit, which in all cases is the 60S subunit (typically 75-90% of the 40S subunit). 60S subunits increased both in absolute terms and per unit cell volume as T cells became activated, reaching a maximum of ∼3.6 million copies per T cell following 2d *in vitro* activation from 264,000 copies in resting T cells (Figure 6F-G). These numbers are similar to those reported by Wolf *et al.* ^42^, but should be more accurate since Wolf *et al.* used total cellular RNA content to estimate ribosomes.

Could there be a significant pool of non-functional ribosomes in the nucleus, where initial assembly occurs, and which occupies nearly 50% of the volume of resting T lymphocytes ^43^ and 34% of activated T cells ^44^? Immunoblotting of fractionated nuclei shows that the distribution of ribosomes in lymphocytes is similar in resting and activated OT-I T cells and HeLa cells, with only a small fraction of total ribosomal subunits detected in nuclear lysates (Supplemental Figure 6).

The ratio of proteins to ribosomes is critical since it dictates the minimal time to replicate the proteome during cell division. This dropped up to 3-fold as T cells became activated (Figure 6H). Since mammalian ribosomes elongate at ∼6 residues per second ^13,36^, we calculated the minimal time for a ribosome to recreate the proteome based on the protein/maximally assembled ribosome ratios, not accounting for protein degradation or secretion.

For HeLa cells, it would take a ribosome 19.9 h to synthesize 910 “average” proteins of 472 amino acids, reasonably close to the reported doubling time of ∼24 hours. For OT-I T cells, with an *in vitro* doubling time of ∼9.7 h, the calculated minimal proteome duplication time is also within shouting distance – 10.1 h by day 2. Therefore, the division rates of *in vitro* activated OT-I T cells, and HeLa cells can be approximated from the number of proteins and functional ribosomes translating a full capacity.

### Paradoxical discrepancy in OT-I cell division rate and protein synthesis capacity

We extended these findings to OT-I T cells *in vivo*, determining first that adoptively transferred OT-I T cells divide most rapidly between day 1 and 2 of activation during acute viral infection, with an average doubling time of 6.8 h, slowing to approximately 7.7 h by day 2 post-infection (via CFSE labeling; Supplemental Figure 3C-E). We sorted for transferred OT-I T cells on day 2 post-infection and measured cell size, protein, and ribosome numbers (“*ex vivo* day 2” measurements in Figure 6 graphs). Cells activated *in vivo* were similar in size and protein content to *in vitro* activated cells, but the protein-to-ribosome ratio was significantly higher than *in vitro* activated T cells due to the presence of 2.3 million *vs.* 3.6 million 60S subunits in maximally *in vitro* activated T cells.

Remarkably, the ratio of proteins to ribosomes (1017) at this juncture dictates a minimal proteome duplication time of 22.2 h, nearly 3x the measured doubling time of 7.7 h. While our RTA measurements support a higher elongation rate *in vivo* (t½=55 sec *vs.* 70 sec in HeLa cells), the 27% increase (7.6 residues per second) does not come close to accounting for the discrepancy. Thus, a paradox: protein synthesis activity or capacity of *in vivo* activated T cells does not support their doubling times.

## Discussion

Lymphocytes protect jawed vertebrates against viral and cellular microbes and tumors. To counter the essentially infinite diversity of antigens expressed by pathogens, lymphocytes evolved to generate an enormous repertoire of specific antigen receptors. Hosts hope to never need the repertoire, and until a cognate antigen appears, metabolic processes, including protein synthesis, are minimized (this might also extend the life span of naive cells since the generation of defective ribosomal products (DRiPs) and other damaging chemical byproducts will also be minimized). Upon activation, lymphocytes divide rapidly to achieve numbers capable of exerting effective immunity. Here, we studied aspects of protein synthesis in lymphocytes, a field fairly dormant since the pioneering studies by the Kay and Cooper laboratories in the 70’s but now experiencing a renaissance ^31,42,45–47^.

To measure translation elongation rates *in vivo* and *in vitro*, we developed flow RTA, which is far simpler and cheaper than original ^36^ and recent ^13,24^ methods and provides information at the level of individual cells simultaneously phenotyped by standard flow cytometry markers. While this work was in progress, the Pierre lab described “SunRiSE”, a nearly identical approach, to measure elongation rates *in vitro*, observing similar puromycylation decay rates following HAR treatment ^27^. While our findings are similar regarding the elongation rates of fibroblasts and lymphocytes, the addition of elongation inhibitors to the protocol (CHX or EME) greatly reduces the leakiness of harringtonine, thereby improving the calculated elongation rate accuracy.

RTA revealed exciting facets of lymphocyte translation. We find that a significant fraction of *ex vivo* lymphocytes possess stalled/primed ribosomes that puromycylate nascent chains but do not transit mRNA. Graber *et al.* ^23^ used the original RPM protocol to show that primary neurons possess substantial numbers of stalled ribosomes, apparently to facilitate rapid translation upon synaptic signaling. Otherwise, to our knowledge, such prolonged ribosomal stalling has not been described in mammalian cells. These experiments may also be useful in examining the phenomenon of “poised mRNA”, originally described in lymphocytes for cytokine mRNAs and more recently expanded on with advanced sequencing techniques ^48,49^.

Our improved RTA protocol reveals the dramatic upregulation of protein synthesis by OT-I CD8^+^ T cells activated *in vivo,* with a 15-fold increase in translation in day 1 activated OT-I T cells *vs.* resting OT-I T cells. Activated T cells divide every 6.8 h from day 1 to day 2 post-VACV-SIINFEKL infection. This is consistent with a prior OT-I study in mice infected with a different SIINFEKL expressing VACV ^50^. Importantly, by i.v. delivery of translation inhibitors, we show that RTA can be used to measure elongation rates *in vivo*. Though we focus on lymphocytes, *in vivo* RTA can be used to study any cell type in animals that can be analyzed *ex vivo* by flow cytometry.

Contrary to observations *in vitro*, ribosomes are not stalled in naïve mouse T cells *in vivo*, as we show via RTA analysis of non-activated T cells. Importantly, ribosome transit times were up to ∼30% faster in activated cells, consistent with the idea that lymphocytes can accelerate translation to support activation and the rapid cell division that ensues. Similarly, OT-I T cells increased elongation rates *in vitro* when incubated at fever temperature (39.5 C°). While such accelerated translation may decrease translational fidelity, the impact may be lessened by the terminal nature of lymphocyte division, since the vast majority of activated cells apoptose within weeks of activation.

We additionally provide initial measurements of numbers of ribosomes and their protein synthesis activity, key values in accounting for the macroeconomics of T cell protein synthesis. Of particular importance is the ratio of cellular proteins to ribosomes; in conjunction with the elongation rate, this value dictates the minimal time (*i.e*., no protein degradation or secretion) for duplicating the proteome ^51^. Using the mammalian cell “speed limit” of 6 residues per second ^13,36^ mouse T cells do not appear to possess sufficient ribosomes to support a 6-8 h division time. Even if ribosomes in *in vivo* activated T cells are translating at 7.8 residues per second, the time required to synthesize the proteome is 2.3x greater than the observed replication time (15.5 h vs. 6.8 h).

The discrepancy is further exacerbated when accounting for protein secretion and degradation of DRiPs (30% of nascent proteins) ^52–55^ and retirees (t½ of ∼32 h over the entire proteome ^56,57^) and the presence of stalled and resting ribosomes. Together, this likely doubles the time required to synthesize the proteome.

Something is obviously wrong. T cell doubling times of 6.8 h are very likely to be accurate, as they are simple to measure and are routinely reported in mouse T cell studies. Quantitating proteins, however, is more challenging than it might seem. Where are the potential gremlins?

1. *Quantitating cellular protein*. We initially used the various dye binding assays for quantitating cellular protein content. While these assays vary notoriously for quantitating different proteins, they provide similar values for the cellular proteome. Moreover, they were in good agreement with a completely independent method based on Trp fluorescence, which we consider the gold standard for protein quantitation ^41^. We use Wisniewski and Gaugaz’s value for Trp abundance in the proteome (0.69%) and note that this value is nearly identical to values obtained using the abundance of Trp toted up from proteomic analysis of 11 different human tumor cell lines ^58^ as well as a transgenic T cell ^59^. One possible source of error is the free metabolic pool of tryptophan, but this is likely to be less than 5% of protein tryptophan ^60^. Another is the presence of serum proteins in cellular lysates. These are unlikely, however, to significantly contribute to our values since we fail to see FBS-derived-BSA in SDS-PAGE of total cell lysates.
2. *Quantitating Ribosomes.* We originally quantitated ribosomes using antibodies specific for ribosomal proteins in immunoblots of total cell lysates with purified ribosomes as a standard. We eventually recognized, however, that this approach is limited by heterogeneity in ribosome composition ^61,62^, as well as the presence of free pools of any given ribosomal subunit. It dawned on us that ribosomes are, in principle, simple to quantitate based on their RNA species, which account for >80% of total cellular RNA. Though we initially quantitated rRNA species on agarose gels with purified ribosomes as a standard, we believe quantitation is more accurate using a Bioanalyzer with doped-in highly purified RNA as an absolute staining standard and a control for yield loss during purification of samples.
3. *Ribosome Elongation Rates.* The classical value of ∼6 or fewer residues per second for radiolabeling studies of cultured cells seems likely to be accurate based on ribosome profiling ^13,63^. A recent ribosome profiling study extends these findings to mice, with *in vivo* elongation rates of 6.8, 5.2, and 4.4 amino acids per second for liver, kidney, and skeletal muscle, respectively ^64^. Our *in vivo* RTA data demonstrate that translation elongation in *in vivo* activated T cells is 30% faster than in cultured cells and thus is likely to be up to ∼7.8 residues/sec. We further note that it is likely that puromycylation detects only a subset of nascent chains. Indeed, in dozens of studies (including our Figure 2C), immunoblots of puromycylated proteins detect discreet bands in SDS-PAGE gel rather than the expected smear if all chains are randomly puromycylated at all lengths. This may be due to non-random incorporation of puromycin, non-random antibody detection of puromycylated nascent chains, or a combination of both. Though it seems improbable, it is possible that this bias influences RTA inferred elongation rates.

While one or more of these values may yet be inaccurate, we note that Wolf *et al.*’s ^42^ mass spectrometric-based measurements of *in vitro* T cell protein synthetic capacity supports and even exacerbates the paradox. We must therefore consider the possibility that that lymphocytes are in such a hurry to divide that they resort to the extraordinary measure of acquiring proteins from resting lymphocytes or other cell types.

There are reports that neurons acquire ribosomes from Schwann cells ^65,66^ and that cancer cells acquire mitochondria from immune cells ^67^. Further, through trogocytosis, lymphocytes acquire cell surface molecules from other cells ^68^. We are, however, proposing that lymphocytes acquire a significant fraction of their proteome, perhaps via something akin to emperipolesis or entosis ^69^, where cells actively enter homotypic cells and can even divide while residing inside ^70^.

In any event, our findings clearly indicate how much remains to be learned about basic lymphocyte cell biology and the importance of simple accounting in squaring our models of cell biology with reality.

## Acknowledgments

We are grateful to Chris Nicchitta (Duke University) for purified ribosomes and Glennys Reynoso for outstanding technical assistance. This work was supported by the Division of Intramural Research, National Institute of Allergy and Infectious Diseases.

## Experimental Procedures

### Mice

Specific-pathogen-free C57BL/6 mice were purchased from the Jackson Laboratory or from Taconic. OT-I TCR transgenic mice were acquired from the NIAID Intramural Research Repository. All mice were housed under specific pathogen-free conditions (including murine norovirus, mouse parvovirus, and mouse hepatitis virus) and maintained on standard rodent chow and water supplied ad libitum. All animal studies were approved by and performed in accordance with the Animal Care and Use Committee of the National Institute of Allergy and Infectious Diseases. For acute infections, and to generate memory T cells, CFSE-labeled Ly5.2+ (CD45.2^+^CD45.1^-^) OT-I T cells were adoptively transferred into Ly5.1 (CD45.1^+^CD45.2^-^) mice. A subset of these mice was infected with VACV-SIINFEKL for indicated times to activate OT-I T cells, with some mice left uninfected where specified. For experiments done directly on memory OT-I T cells, assays were done 8-9 weeks after infection.

### *In vivo* RPM, *in vivo* RPM-RTA, and relative protein synthesis determination

For the standard and CHX-improved *in vivo* RPM assays, mice were intravenously injected with 100μl of a 10mg/ml solution of PMY in PBS that was warmed to 37°C, or PMY, as just described, along with 0.34mg per mouse of CHX. After indicated times, mice were sacrificed, and organs were collected into complete RPMI on ice (Gibco RPMI supplemented with 7.5% fetal calf serum). For the *in vivo* RTA, or the CHX-improved *in vivo* RTA, mice were intravenously injected with 100μg of HAR simultaneously with 1mg of PMY, or 1mg of PMY and 0.34mg of CHX for 5 minutes (for the maximum signal) or first intravenously injected with 100μg of HAR for the times indicated before being intravenously injected with 1mg of PMY, or PMY and 0.34mg of CHX for 5 minutes. To determine relative levels of active protein synthesis, two sets of mice were required. In the first set, mice were intravenously injected with 100μg of HAR for 15 minutes, and then intravenously injected with 1mg of PMY and 0.34mg of CHX for 5 minutes before spleens were harvested. In the second set, mice were intravenously injected simultaneously with 0.34mg of cycloheximide and 1mg of PMY for 5 minutes before spleens were harvested.

### Single cell preparation from organs

Isolated organs were crushed between two frosted microscope slides, and the resultant single cell suspension was filtered through a 70µm mesh screen. The filtered single cell suspension was then centrifuged, resuspended in ACK lysing buffer (Lonza) to lyse red blood cells, centrifuged again, and resuspended in complete RPMI for counting on a Nexcelom Cellometer Vision using Trypan Blue (Lonza BioWhittaker) for live/dead cell discrimination and cell diameter measurements.

### CFSE labeling

Spleens and inguinal, mediastinal, cervical, mesenteric, and popliteal lymph nodes from OT-I TCR transgenic or C57BL/6 mice were processed into a single cell suspension, red blood cells were lysed in ACK lysing buffer, and the resultant cells filtered through a 70 µM mesh screen. After two washes in PBS, cells were counted on a Nexcelom Cellometer Vision using Trypan Blue for dead cell exclusion, and cells were labeled in 5 µM CFSE (Invitrogen) in PBS at 1×10^7^ cells per ml for 18 minutes in a 37° water bath with mixing every 6 minutes. Cells were washed three times in PBS, recounted, and adoptively transferred into the indicated mice or cultured as specified.

### Human lymphocyte purification and culture conditions for human and mouse lymphocytes

Elutriated human lymphocytes were from healthy anonymous donors at the NIH Clinical Center Department of Transfusion Medicine. After collection, elutriated lymphocytes were purified on a discontinuous 35-70% Percoll (Amersham Biosciences) gradient and washed once with ACK lysing buffer (Life Technologies) to remove contaminating red blood cells. For time-course experiments, purified lymphocytes were resuspended in PBS and labeled with CFSE where indicated (as described above) to enable tracking of cell division over time. Lymphocytes were plated at 1-2 × 10^6^ cells/mL in RPMI: RPMI 1640 (Gibco) supplemented with 15% FCS, 25mM HEPES (Corning Cellgro), 1mM sodium pyruvate (Gibco), and 55uM BME (Gibco). Depending on the experiment, media was also supplemented with recombinant human IL-2 (BRB NCI Frederick, 25 U/mL), PMA (Sigma, 1ng/mL), and ionomycin (Sigma, 100ng/mL). For OT-I T cell cultures, PMA was added at 100ng/ml instead, and, where noted, SIINFEKL was added as well (100nM) for optimal activation. Lymphocytes were cultured in 6% CO2 at 37°C and allowed to sit overnight prior to any experiments unless noted (noted as “freshly isolated”). For time-course experiments, lymphocytes were cultured for up to 5 days and resuspended in fresh media every 2 days. Cell counts, diameters, and viabilities (through Trypan blue exclusion) were made on a Nexcelom Cellometer Vision cell counter. Cell volumes were calculated assuming spherical geometry.

### *In vitro* RPM and RPM staining

For each sample, cells were resuspended at 2 × 10^7^ cells per mL and 100μl transferred into 96-well round-bottom plates. When indicated, the media contained protein synthesis inhibitors at the following concentrations: 5μg/mL HAR (Santa Cruz Biotechnology), 25μg/mL EME (Calbiochem), 200μg/mL CHX (Sigma), 50μg/mL anisomycin (Sigma), or 10μM pactamycin (Sigma). After a 15-minute incubation at 37°C, 50μL of 3X PMY (Calbiochem) media was added (150μg/mL, for a final concentration of 50μg/mL) and the cells were incubated for an additional 5 minutes before shifting to ice and adding 100μL of cold PBS. Cells were then stained with ethidium monoazide (10μg/mL in PBS, Molecular Probes) for live/dead cell discrimination. After thorough washing, and a 10-minute incubation with heat-inactivated sera, or 2.4G2 to block Fc receptors, cell surface antigens were labeled for 30 minutes at 4°C with the following antibodies: For human lymphocyte stains, antibodies against: CD3ε PerCP-eFluor 710 (clone OKT3, eBioscience), CD19 PE (clone HIB19, eBioscience), CD45 APC-eFluor 780 (clone 2D1, eBioscience), CD4 PE-CF594 (clone RPA-T4, BD) or CD4 PE-Cy7 (clone RPA-T4, eBioscience), and CD8α BV421 (clone RPA-T8, BD). For mouse lymphocyte stains, antibodies were: CD3ε BV786 (clone 145-2C11, BD), CD4BV510 (clone RM4-5, BD), CD5 APC-R700 (clone 53-7.3, BD), CD8α PE-CF594 (clone 53-6.7, BD), CD11b PE-Cy7 (clone M1/70, eBioscience), CD19 APC-Cy7 (eBio1D3, eBioscience), CD25 BV650 (PC61, BD), CD44 BV605 (IM7, BD), CD44 eFl450 (clone IM7, eBioscience), CD45.1 APC (clone A20, eBioscience), CD45.1 eFl450 (clone A20, eBioscience), CD45.2 eFluor450 (clone 104, eBioscience), CD45.2 PE-Cy7 (clone 104, eBioscience), CD69 PerCP-Cy5.5 (clone H1.2F3, Invitrogen), γδ TCR PE (eBioGL3, GL3, eBioscience), Gr1 PE (clone RB6-8C5, BD), NK1.1 FITC (clone PK136, eBioscience), TCRβ 711 (H57-597, BD), and Vβ5.1/Vβ5.2 PE (clone MR9-4, BD). All antibodies were used at 1:150 dilution in buffered saline supplemented with 0.1% BSA. Next, cells were simultaneously fixed and permeabilized in fix/perm buffer (1% PFA, 0.0075% digitonin in PBS) for 20 minutes at 4°C. Intracellular PMY was labeled with an anti-PMY antibody (clone 2A4) directly conjugated with Alex Fluor 647 (conjugated using the Life Technologies Protein Labeling Kit per the manufacturer’s instructions) for 1 hour. Cells were thoroughly washed and resuspended in buffered saline supplemented with 0.1% BSA, flow cytometry performed on a BD LSRII or BD LSRFortessa X-20, and resulting data analyzed with FlowJo software. To gate on OT-I T CD8^+^ T cells, setup was: singlets by FSCa and FSCw, lymphocytes by SSCa and FSCa, EMA^-^ (live/dead cell marker), CD3^+^CD19^-^, CD8^+^CD4^-^, CD45.2^+^CD45.1^-^, and Vb5^+^, and activation markers as indicated. For thymocyte subsets, gating setup was singlets by FSCa and FSCw, lymphocytes by SSCa and FSCa, EMA^-^, and then subsets on combinations of CD3ε, CD4, CD8α, CD19, CD25, CD44, CD69, γδ TCR, and TCRβ. For human lymphocytes, gating setup was singlets by FSCa and FSCw, lymphocytes by SSCa and FSCa, EMA-, and on subsets as indicated.

### Amino acid radiolabeling

The following reagents were used for radioactive amino acid labeling: DMEM (for labeling HeLa cells) or RPMI minus leucine (RPMI without L-leucine, L-glutamine, and sodium pyruvate from MP Biomedicals, supplemented with Glutamax and 1mM sodium pyruvate) for labeling human lymphocytes with or without inhibitors and for the [^3^H]-Leu (Perkin Elmer) ribosome transit analysis. Cells were kept at 37°C throughout the experiment and labeling. Cells were resuspended in complete RPMI at 1 × 10^7^ cells/ml and 1ml was aliquoted into fresh Eppendorf tubes. Next, cells were spun at 300g for 4 minutes and pre-treated with protein synthesis inhibitors (concentrations as in RPM Staining above) in complete RPMI for 15 minutes. Pre-treated cells were spun at 300g for 4 minutes, resuspended in 200μL of labeling media (RPMI-Leu) containing 0.2mCi/mL [^3^H]-Leu in the absence or presence of protein synthesis inhibitors as indicated. After a 5-minute labeling period, protein synthesis was stopped by adding 1mL of ice-cold PBS containing 200μg/mL of CHX. For all labeling experiments, after washing cells thoroughly in ice cold PBS, cells were lysed in 100μL or 200μl of 2% SDS lysis buffer (2% SDS, 50mM Tris-HCl pH 7.5, 5mM EDTA, 15U/mL DnaseI (Roche), cOmplete mini EDTA-free protease inhibitor tablet (Roche) in water) and boiled for 30-60 minutes to ensure complete lysis. Protein amounts were quantified by the DC Protein Assay (BioRad) or by tryptophan fluorescence measurements ^41^. To quantify the amount of [^3^H]-Leu incorporated into proteins, equal amounts of lysate (six replicates per condition) were spotted onto a 96-well DEAE filter mat (PerkinElmer) and the mat was dried at 60°C. The mat was then soaked in a 10% trichloroacetic (TCA) (Calbiochem) solution for 30 minutes at room temperature, washed twice in 70% ethanol, dried at 60°C, placed in a MicroBeta sample bag (PerkinElmer) with ∼6mL of BetaPlate Scint (PerkinElmer), and heat sealed. Radioactivity was quantified in a 1450 MicroBeta TriLux scintillation counter. To determine the total amount of amino acid incorporated into proteins, dilutions of the [^3^H]-Leu stock were counted and used as standards.

### *In vitro* RPM Ribosome Transit Analysis (RTA)

RPM-RTA: For each time point, 1 × 10^6^ lymphocytes were transferred to a fresh conical tube and resuspended in 250μL of the appropriate media. Cells were kept at 37°C (or 39.5°C when indicated) throughout the experiment. An equal volume of 2X inhibitor media was added to each tube at the indicated time and the tube was vortexed briefly to mix. Depending on the time course, the 2X inhibitor media contained HAR (Santa Cruz Biotechnology) at 10μg/mL (final concentration 5μg/mL), or HAR at 10μg/mL and EME (Calbiochem) at 50μg/mL (final concentrations of 5μg/mL and 25μg/mL, respectively. At the end of the time course, an equal volume (250μL) of 3X puromycin (PMY) (Calbiochem) media (150μg/mL, for a final concentration of 50μg/mL) was added to each tube and the tube was vortexed briefly to mix. Cells were incubated for 5 minutes with PMY before adding an excess of ice-cold PBS to quench the ribopuromycylation reaction. The cells were then stained for analysis by flow cytometry as described above.

[^3^H]-Leu RTA: Cells were kept at 37°C throughout the experiment. For each time point, 30 × 10^6^ lymphocytes were transferred to fresh Eppendorf tubes and resuspended in 50μL of the RPMI-Leu. 50μl of 2X inhibitor media was added to each tube and the tubes were vortexed briefly to mix. The inhibitor concentrations are as noted above. At the indicated times, an equal volume (100μL) of [^3^H]-Leu labeling media (RPMI-Leu media and [^3^H]-Leu in a 1:1 ratio, 0.5mCi/mL) was added and cells were labeled for 5 minutes. To stop [^3^H]-Leu incorporation, an excess of ice-cold PBS containing 200μg/mL CHX and 1mg/mL leucine was added before placing the cells on ice. Cells were lysed and [^3^H]-Leu incorporation quantified as described under “Amino acid radiolabeling” subsection.

### Polysome profiling

A 15-45% continuous sucrose gradient was made in Thinwall polyallomer tubes (Beckman Coulter) from 15% and 45% sucrose (MP Biomedicals) solutions in gradient buffer (20mM Tris-HCl pH 7.4, 5mM MgCl2, 100mM KCl, supplemented with 100μg/mL CHX (Sigma) and 10U/mL RNaseOUT (Invitrogen)). Briefly, 5mL of the 15% sucrose solution was carefully layered onto 5mL of the 45% sucrose solution, and the tube was placed horizontally at 4°C, typically overnight or for at least 2.5 hours before the experiment. For each gradient, cells were harvested and washed in cold PBS as described above. For cell lysis, cells were first swelled by adding 950μL of a cold hypotonic buffer (20mM Tris-HCl pH 7.4, 5mM MgCl2, 10mM KCl, 40U/mL RNaseOUT, 0.1U/μL SuperaseIn, supplemented with Complete EDTA-free protease inhibitors (Roche)). After 10 min of cell swelling, NP-40 was added to a final concentration of 0.5%, the resultant lysate mixed, incubated on ice for 3 min, and spun at 7000 rpm for 2 min to remove nuclei. Post-nuclear lysates were then brought to 100mM KCl (or 300mM or 500mM NaCl where indicated), layered onto 15-45% continuous sucrose gradients, and spun for 100 min at 38,000rpm at 4°C in a SW41Ti rotor (Beckman Ultracentrifuge). Gradients were syringe-fractionated mechanically from the bottom up and monitored for absorbance at 254nm (Teledyne Isco) to obtain polysome profiles. When indicated, area under the curve measurements were calculated by a trapezoidal method from the resulting curves. When required by the experiment, we concentrated monosome and polysome fractions for immunoblotting. To pellet the ribosomes, we spun the pooled monosome or polysome fractions for 1 hour or O/N at 39,000 rpm at 4°C in the SW41Ti rotor on a 34% sucrose cushion. The resultant ribosome pellet was resuspended in 2% SDS extraction buffer.

### Quantitating cellular proteins

We quantitated total protein in cell lysates based on Trp fluorescence ^41^. Briefly, cells (1-2 million lymphocytes per 100µl) were lysed for 10-30 minutes at 95°C in a solution of 2% SDS, 0.1 M Tris-HCl, 50 mM DTT (pH 7.8) with 15U/mL DnaseI (Roche) and a cOmplete mini EDTA-free protease inhibitor tablet (Roche) added fresh. An 8M urea, 10mM Tris-HCL with 0.5mM DTT solution was freshly prepared, and 200µl added to wells of a flat-bottomed black polystyrene plate, and 2-4 µl of either cell lysates or a Trp standard solution was added to individual wells in triplicate. Fluorescence emitted at 350nm after excitation at 295nm was measured. We also compared this assay with the commercially available DC protein assay (Bio-Rad, performed according to the manufacturer’s instructions), and found that the assays generated similar values.

### Quantitating cellular ribosomes

After lymphocyte isolation, the PBS-washed cell pellet was dissolved in TRIzol; a spike-in mRNA standard was added at this step (CleanCap EGFP mRNA from TriLink, L-7601) to account for RNA loss during processing. RNA purification was conducted as described in the manufacturer’s TRIzol protocol, with 5µg of glycogen used as carrier and the isopropanol precipitation step conducted overnight at -20°C. The final RNA pellet was dissolved in 50µL of ultra-pure water and roughly quantified to determine appropriate range for the Agilent bioanalyzer chip. Samples, including fresh spike-in mRNA alone, were loaded and run on RNA Nano Bioanalyzer chips (Agilent RNA Nano 6000), with a 70°C heating step and run on a 2100 Agilent Bioanalyzer. Bioanalyzer 2100 Expert software was used to determine total RNA concentration of each sample and percent area under curve of each peak (mRNA spike-in at approximately 1000 bp, 18S rRNA at around 1800 bp, 28S rRNA at around 4000 bp). The yield of the RNA prep was calculated as follows:

(mRNA spike-in standard peak from an RNA-purified sample) / (average of 2-3 standard peaks from 75ng/µL standard wells) = (fraction of RNA that remains after the purification process). We next converted the 18S and 28S ng/µL values to ‘number molecules per cell’ using the number of cells that originally went into the RNA purification, the RNA yield described above, and the following values: mouse 18S = 6.40E+05 g/mol; mouse 28S = 1.60E+06 g/mol; human 18S = 6.40E+05 g/mol; human 28S = 1.70E+06 g/mol.

### Immunoblotting

To fractionate cells into nuclear and non-nuclear lysates, cells were either dissolved directly in 2% SDS extraction buffer at 95°C (“all” in Supplemental Figure 6) or subjected to a hypotonic lysis procedure. Cells were swelled with a buffer containing 20mM Tris-HCl pH 7.4, 2.5mM MgCl2, and 10mM KCl supplemented with protease inhibitors for 10 min on ice. NP-40 was added to a final concentration of 0.5%, and the resultant lysate was mixed, incubated on ice for 3 minutes, and spun at 7000 rpm for 1 min. Non-nuclear lysates were removed and immediately dissolved in gel loading sample buffer (Life Technologies) to prevent sample degradation. Nuclei were washed gently 2x with PBS buffer containing NP-40 and protease inhibitors. Finally, nuclear proteins were extracted by dissolving pelleted nuclei in 2% SDS extraction buffer at 95°C. Equal amounts of each fraction were prepared for SDS-PAGE.

Samples were electrophoresed in 4-12% NuPAGE Bis-Tris gels (Invitrogen). Proteins were then transferred to nitrocellulose membranes (iBlot system, Novex) and membranes stained with Ponceau S and washed with PBS to confirm transfer uniformity. Next, membranes were incubated with either StartingBlock buffer (ThermoScientific) or Odyssey Blocking Buffer (Licor), followed by primary antibody prepared in StartingBlock buffer or Odyssey Blocking Buffer with 0.1% Tween-20 (Sigma). Depending on the experiment, we used the following primary antibodies: mouse anti-PMY (clone 2A4) at 6.66μg/mL; human anti-ribosomal P antigen at 1:2000 (Immunovision); rabbit anti-RPL7 at 1:1000, rabbit anti-RPL26 at 1:2000 (Bethyl); mouse anti-beta actin at 1:4000 (Licor); rabbit anti-HSP90 at 1:500 (Santa Cruz); rat anti-GRP94 at 1:1500 (Novus); rabbit anti-RPL28 at 1:500, rabbit anti-RPL6 at 1:1000, mouse anti-PDI at 1:2000 (Abcam); mouse anti-lamin A/C at 1:2000, rabbit anti-fibrillarin at 1:1000, mouse anti-RPS6 at 1:1000, rabbit anti-histone H3 at 1:2000, rabbit anti-RPL5 at 1:1000, and rabbit anti-RPS3 at 1:1000 (Cell Signaling Technology). The number of ribosomes per cell in earlier experiments was quantified by generating a standard curve using highly purified HeLa cell or canine rough microsome ribosomes (a kind gift of Chris Nicchitta, Duke University).

Membranes were washed three times in PBS + 0.1% Tween-20 (PBS-T) followed by incubation with secondary antibodies (all from Licor; used at 1:10,000) prepared in StartingBlock buffer or Odyssey Blocking Buffer. Membranes were washed 3x in PBS-T, 1x in PBS, and scanned via a Licor Odyssey CLX scanner.

---

**Supplemental Figure 1 – RPM tracks translation in distinct cell populations over time**

(A) Population frequency and RPM of resting day 2 or day 5 human lymphocytes or PMA/ionomycin/IL-2 activated day 2 or day 5 human lymphocytes. Left panel is the percent of CD45^+^ cells in the indicated population, the right panel is the RPM signal in each population. (B) Representative RPM flow cytometry plot gated on polyclonal CD8^+^ T cells. Similar data was obtained from all donors. (C) Gating strategy to quantify HeLa cell RTA assays as described in Figure 2.

**Supplemental Figure 2 – Dominant populations of monosomes in resting human and mouse lymphocytes**

(A) Primary human lymphocytes were cultured for one day in the absence of PMA/ionomycin or for up to 2 days in the presence of PMA/ionomycin/IL-2, followed by polysome profiling. Representative of two independent experiments. (B) Freshly isolated human lymphocytes were treated with 0.1μg/mL CHX for 30 minutes prior to cell lysis and sucrose gradient centrifugation. Representative of two independent experiments. (C) C57BL/6 mice were treated IV with vehicle or CHX. After 10 min, spleens and lymph nodes were harvested, and the resulting cells subjected to polysome profiling via ultracentrifugation through 15-45% sucrose gradients. (D) Lymphocytes or hepatocytes were harvested from OT-I mice treated IV with CHX, lysed, and processed for polysome profiling. For activated cells, OT-I T cells were treated with PMA/ionomycin and IL-2 for 2 days prior to CHX treatment and polysome profiling. Bottom right; quantification of the areas under the curve of free subunits, monosomes, and polysomes. Representative of two independent experiments.

**Supplemental Figure 3 – RPM cell phenotyping and *in vivo* T cell division**

(A) C57BL/6 mice were treated intravenously with CHX and PMY or only PMY. After the indicated times, splenocytes were harvested, surface stained, fixed/permeabilized, and RPM staining was performed. Representative of three independent experiments, 2-3 mice per group. (B) In one set of C57BL/6 mice, HAR was intravenously injected for 15 minutes before intravenously injecting mice with PMY for 5 min. In a second set of mice, CHX and PMY were IV injected for 5 min. Splenocytes from each set of mice were harvested, surface stained, fixed, and permeabilized, and RPM staining was performed for various immune cell subsets. To determine relative amounts of ribosomes, the ‘HAR then PMY’ RPM signal was subtracted from the CHX+PMY RPM signal for each cell subset after flow cytometry. Representative of two independent experiments, 2-4 mice per group. (C) CFSE-labeled Ly5.2^+^ (CD45.2^+^CD45.1^-^) OT-I cells were adoptively transferred into Ly5.1^+^ (CD45.1^+^CD45.2-) mice, which were then infected with VAC-SIINFEKL. 1-3 days after infection, mice were intravenously injected with CHX simultaneously with PMY for 5 min. Splenocytes from the mice were harvested, surface stained, fixed and permeabilized, and RPM staining was performed. Representative flow cytometry plots gated on OT-I T cells. (D) Gating strategy used to display and quantify *in vivo* T cell data as described in panel C and elsewhere. (E) Number of divisions (by CFSE dilution) of OT-I T cells 1-3 days after infection of mice with VACV-SIINFEKL. (F) Amount of translation as measured by RPM signal (with “no PMY” signal subtracted) in uninfected, or one-, two-, or three-day VACV-SIINFEKL-infected mice. Representative of four independent experiments, 2-3 mice per time point.

**Supplemental Figure 4 – Exogenous SIINFEKL significantly enhances OT-I T cell activation *in vitro***

(A) Maximum-normalized MFI of CD69, CD25, CD44, and side scatter (SSCa), as well as CFSE expansion index, of OT-I T cells after 1 or 2 days with PMA/ionomycin or PMA/ionomycin with exogenous SIINFEKL. IL-2 was included in all conditions. (B) Side scatter is a good proxy for cell size. SSCa MFI plotted vs cell diameter as determined by automated cell counter measurements.

**Supplemental Figure 5 – Salt stability of T cell ribosomes and monosome quantification**

(A) Lymph node and splenic OT-I T cells were mixed and stimulated *in vitro* for 2 days with SIINFEKL, PMA/ionomycin, and IL-2. Cells were treated with CHX for 5 minutes, lysed, and brought to either 300 or 500mM NaCl final concentration. Ribosome-containing lysates were subjected to polysome profiling via ultracentrifugation through 15-45% sucrose gradients containing either 300 or 500mM NaCl. (B) Quantification of the areas under the curve of free subunits, monosomes, and polysomes for each sample. (C) For resting T cells, OT-I mice were treated IV with CHX and lymphocytes from the spleens or lymph nodes were isolated and lysed. For activated T cells, lymph node and splenic OT-I T cells were mixed and stimulated *in vitro* for 2 days with SIINFEKL, PMA/ionomycin, and IL-2, treated with CHX for 5 minutes, and lysed. Lysates were subjected to polysome profiling via ultracentrifugation on 15-45% sucrose gradients after bringing both lysate and sucrose gradients to a final concentration of 500mM NaCl to dissociate non-translating ribosomes. (D) Quantification of the areas under the curve of free subunits, monosomes, and polysomes in each sample. Representative of two independent experiments.

**Supplemental Figure 6 – Fractionation of HeLa or T cells reveals few ribosomal components in nuclear lysates**

HeLa cells, freshly isolated resting OT-I T cells, or OT-I T cells stimulated with PMA/ionomycin and IL-2 *in vitro* for 2 days were either lysed directly in SDS extraction buffer (“all”) or subjected to a hypotonic lysis procedure to isolate non-nuclear lysates and nuclear lysates. Equal amounts of each fraction were subjected to immunoblotting for markers typical of the cytosol, ER, and nucleus. Antibodies against ribosomal proteins were used to determine where the majority of ribosomal proteins (and therefore ribosomes) fractionated. Controls with antibodies specific for nucleolar located fibrillarin, histone H3, and lamin A/C establish lack of nuclear contamination in non-nuclear fractions. ER and cytoplasmic proteins HSP90, GRP94, PDI, and actin indicate lack of contamination in the nuclear fraction.

