## Supplementary figures and images for "Paradox Found: Global Accounting of Lymphocyte Protein Synthesis"

### Supplemental Figures

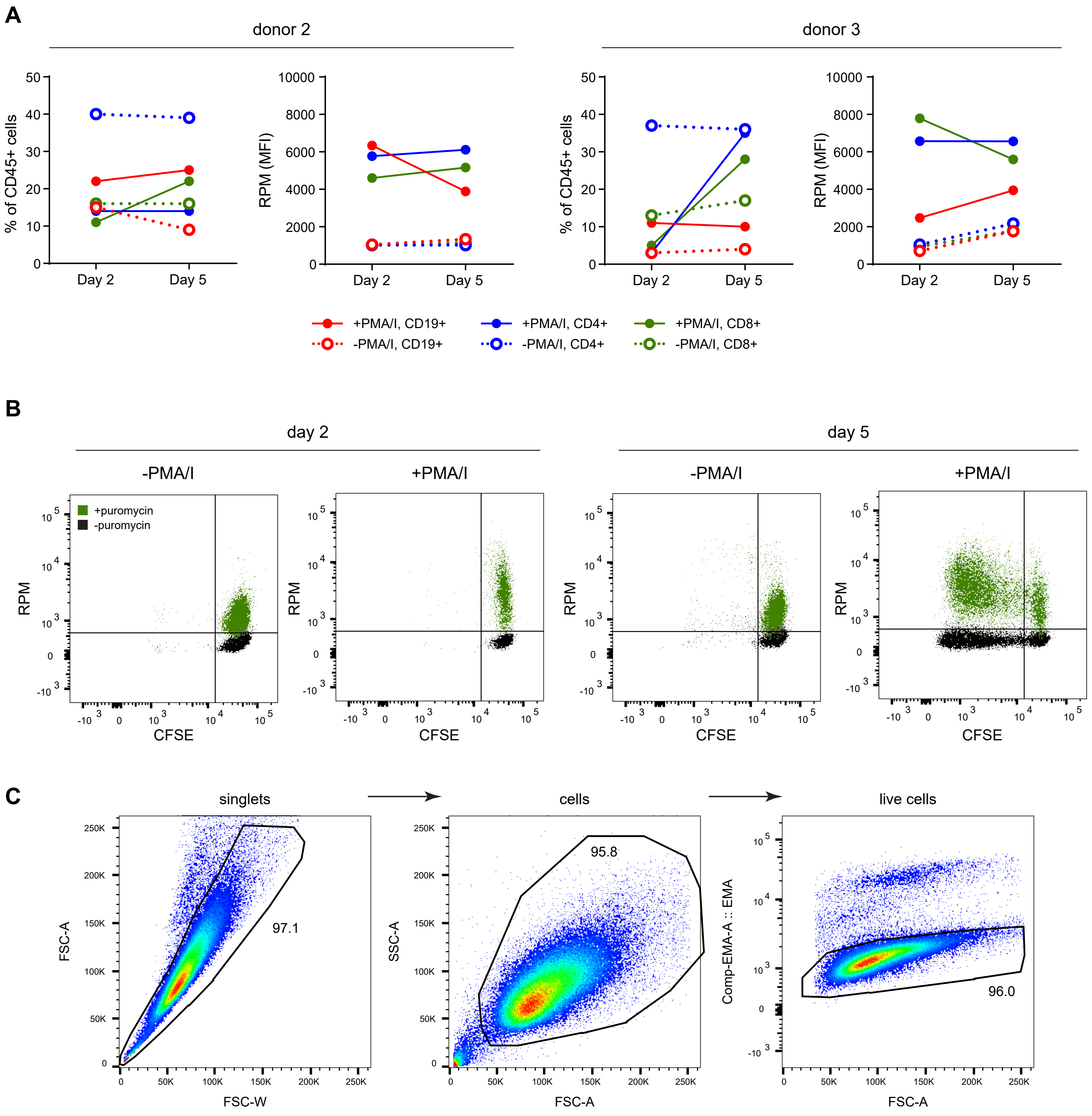

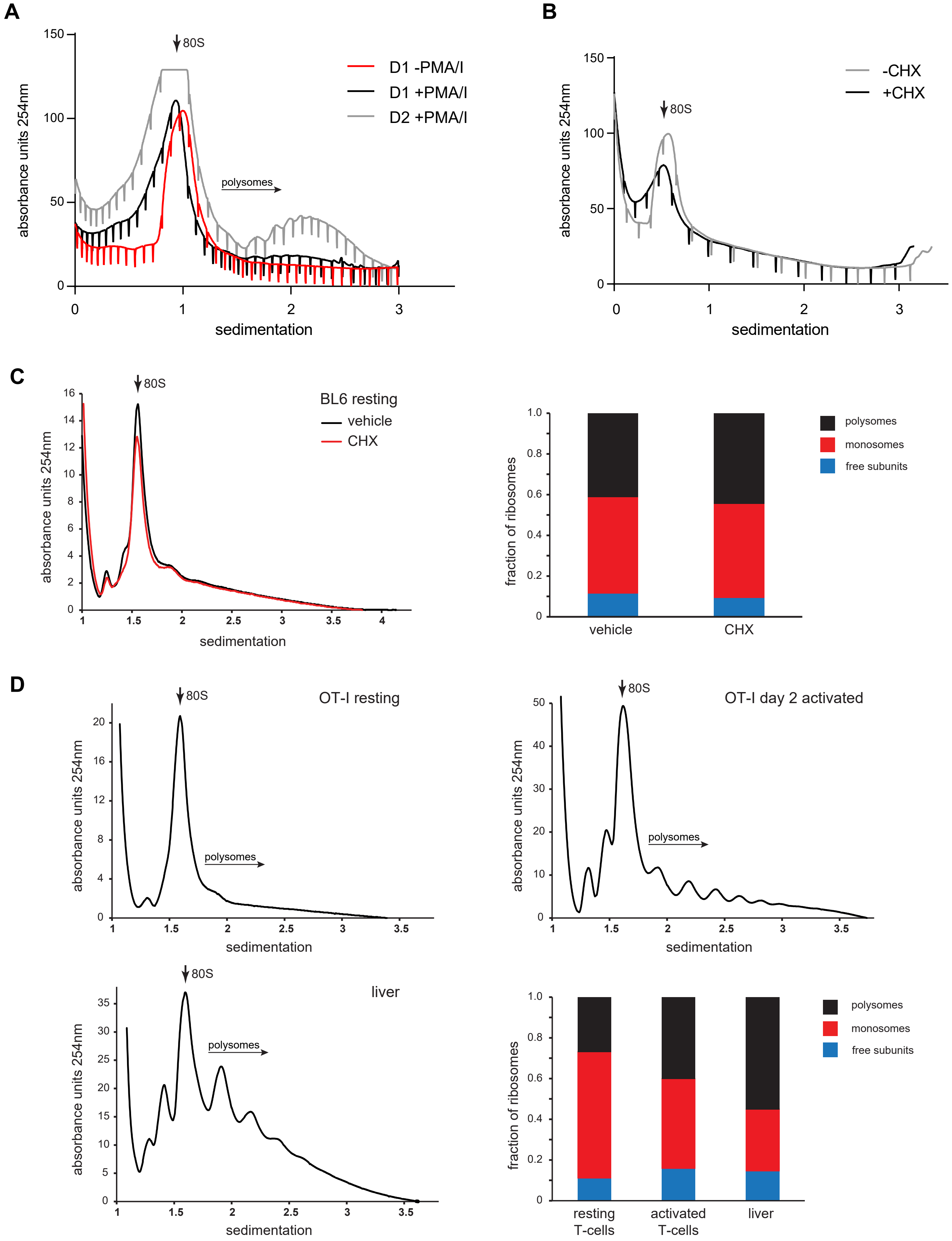

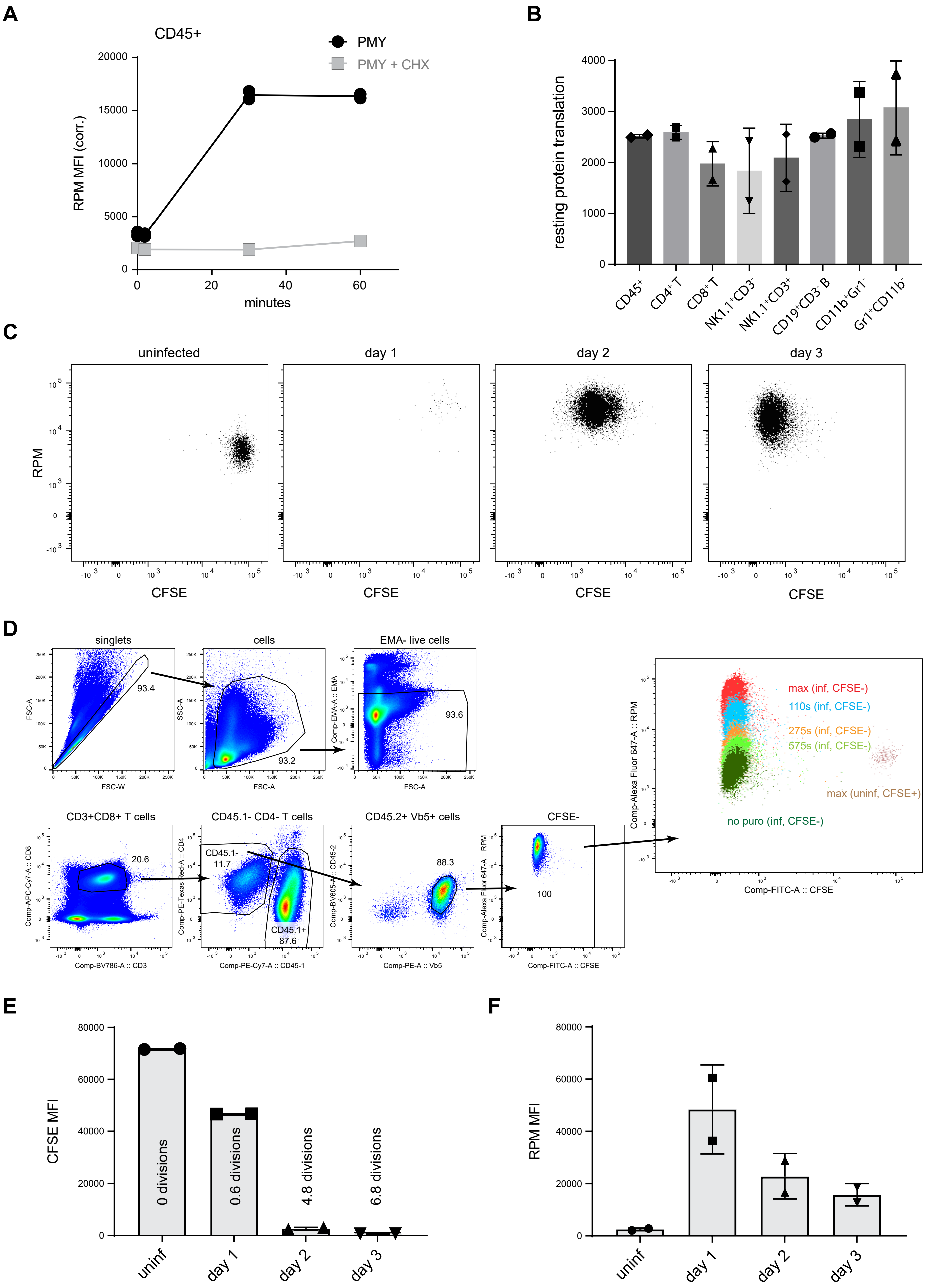

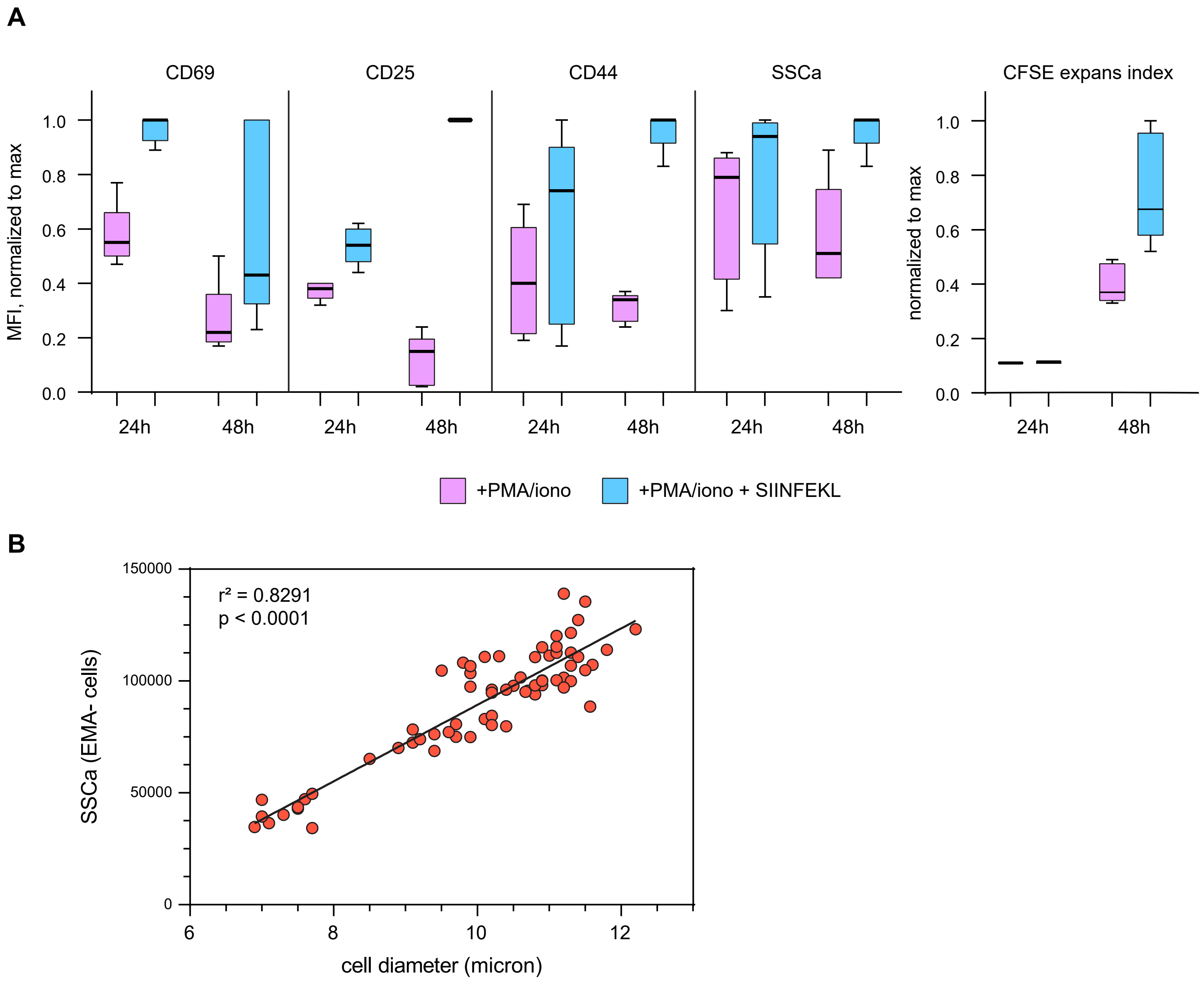

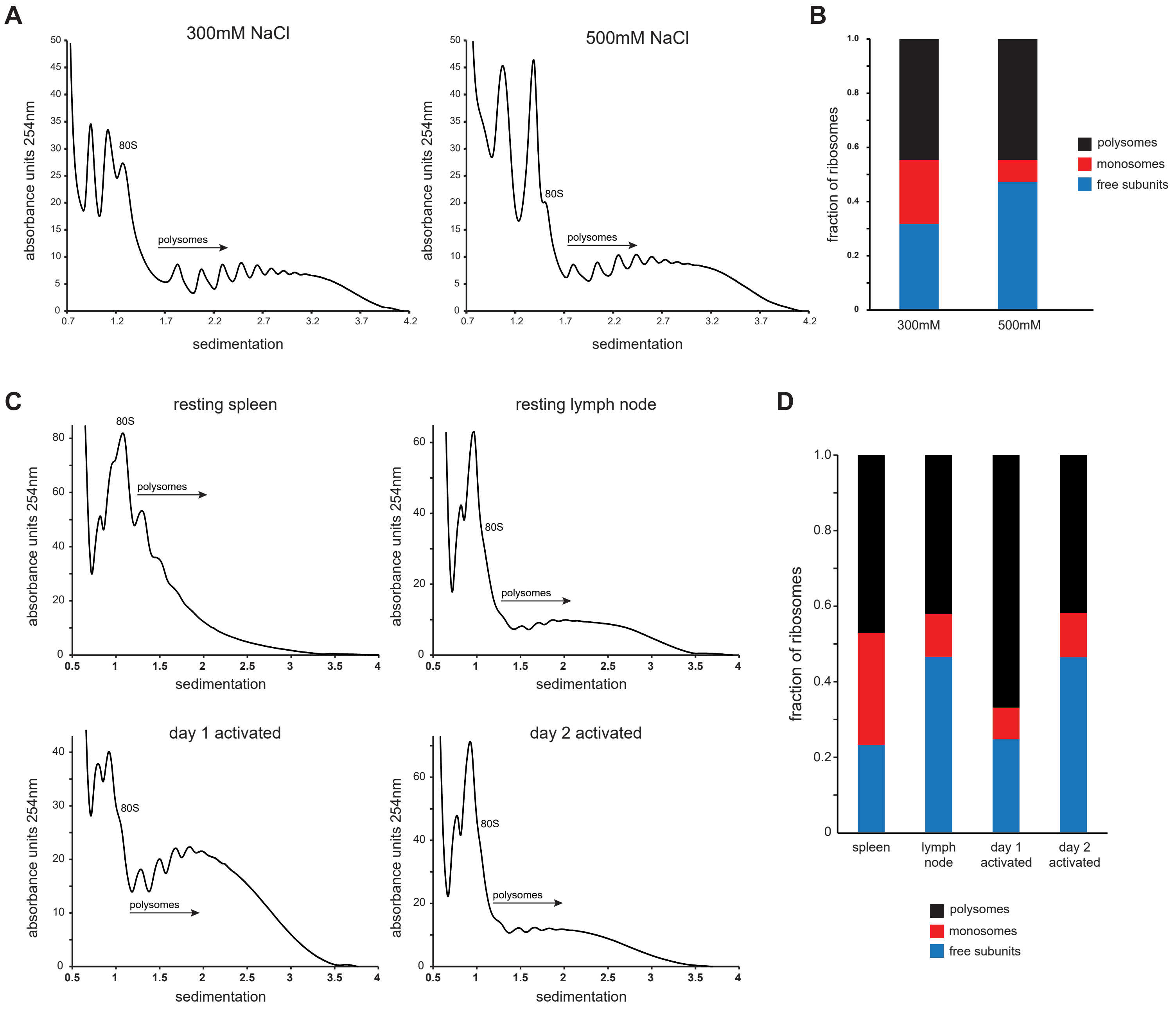

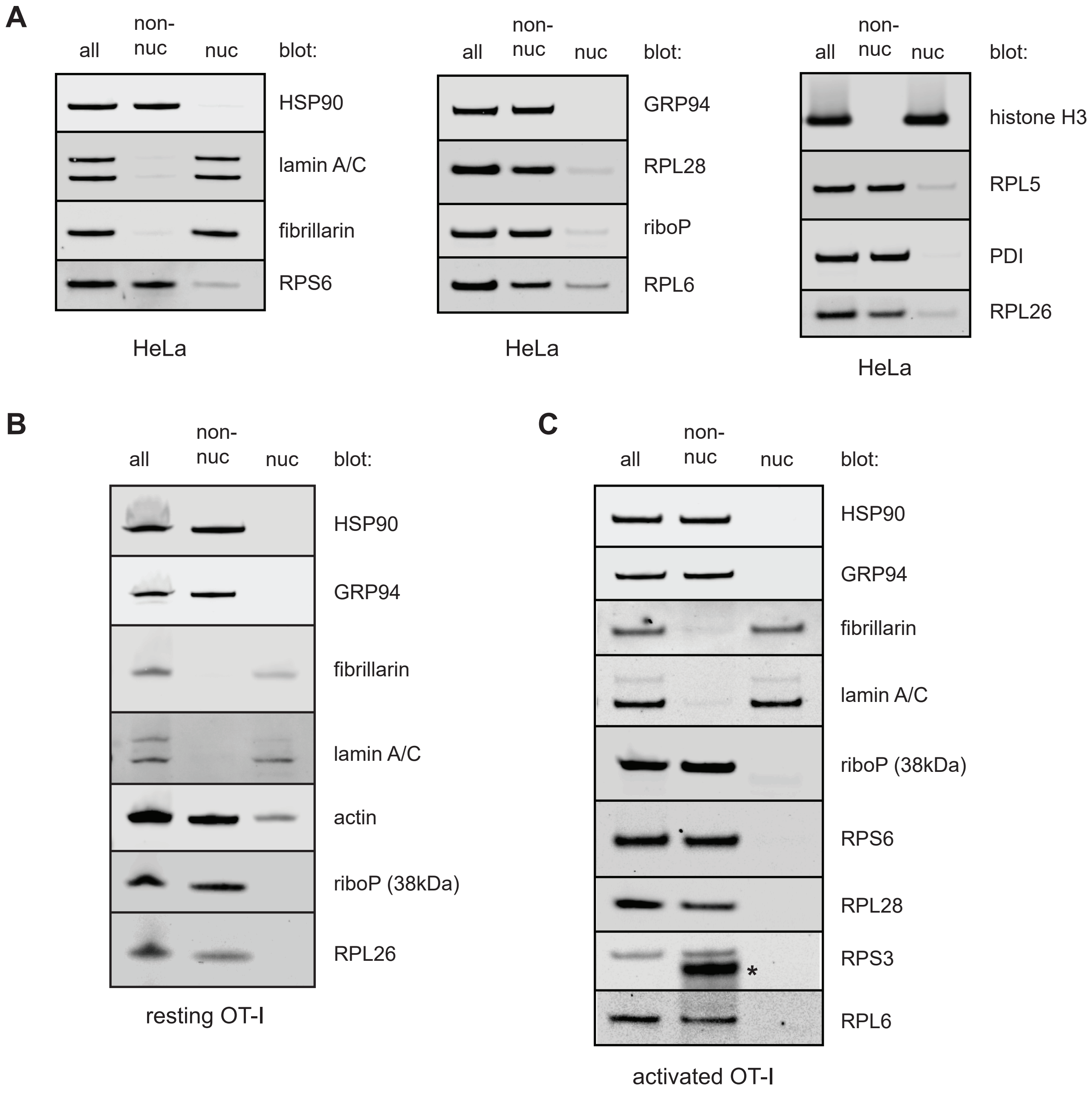
